# A Tunable, Ultrasensitive Threshold in Enzymatic Activity Governs the DNA Methylation Landscape

**DOI:** 10.1101/2024.06.25.600710

**Authors:** Kwadwo A. Bonsu, Annie Trinh, Timothy L. Downing, Elizabeth L. Read

## Abstract

DNA methylation is a widely studied epigenetic mark, affecting gene expression and cellular function at multiple levels. DNA methylation in the mammalian genome occurs primarily at cytosine-phosphate-guanine (CpG) dinucleotides, and patterning of the methylation landscape (i.e., the presence or absence of CpG methylation at a given genomic location) exhibits a generally bimodal distribution. Although much is known about the enzymatic writers and erasers of CpG methylation, it is not fully understood how these enzymes, along with genetic, chromatin, and regulatory factors, control the genome wide methylation landscape. In this study, methylation is analyzed at annotated CpG Islands (CGIs) and independent CpGs as a function of their proximity to other CpG substrates. Analysis is aided by a computationally efficient stochastic mathematical model of methylation dynamics, enabling parameterization from data. We find that methylation exhibits a switch-like dependence on local CpG density. The threshold and steepness of the switch is modified in cell lines in which key enzymes are knocked out. Modeling further elucidates how enzymatic parameters, including catalytic rates and lengthscales of inter-CpG interaction, tune the properties of the switch. Together, the results support a model in which competition between opposing TET1-3 demethylating enzymes and DNA methyltransferases (DNMT3A/B) results in an ultrasensitive switch, analogous to the protein phosphorylation switch (termed ‘zero-order ultra-sensitivity’) proposed by Goldbeter and Koshland. Our study provides insight to the mechanisms underlying establishment and maintenance of bimodal DNA methylation landscapes, and further provides a flexible pipeline for gleaning molecular insights to the cellular methylation machinery across cell-specific, epigenomic datasets.

## Introduction

DNA is organized into chromatin structures that define its state and work in concert to regulate activation and silencing of the genome. Epigenetic modifications that modify chromatin and modulate gene expression include (i) non-coding RNAs, (ii) covalent histone modifications, (iii) positioning of nucleosomes and incorporation of histone variants and (iv) methylation of cytosine residues. These components synergize to control expression of the genome (1, 2).

Of these epigenetic mechanisms, DNA methylation is one of the most widely conserved amongst mammals, and as such has been broadly studied. In the mammalian genome, methylation occurs typically at cytosine-phosphate-guanine (CpG) dinucleotides at the 5th carbon position of cytosine. CpGs are known to cluster at certain loci, termed “CpG Islands” (CGIs), with highly variable sizes across the human genome (3–5). Promoter regions are frequently associated with CGIs of various size and CpG content, and CGI methylation is anti-correlated to promoter accessibility/activation (2, 6–8). There is also high CpG content associated with Transposable Elements (TEs) such as Short- and Long-Interspersed Retro-transposable Elements, SINEs and LINES respectively, and Long-Terminal Repeats (LTRs). These regions are predominantly methylated and their ability to disrupt normal regulation is suppressed (9).

Catalytically-active DNA Methyltransferases (DNMTs) are responsible for deposition of methyl marks, and they actively contribute to methylation patterning during replication, development, and in disease. DNMT1 is primarily responsible for maintenance methylation, or re-establishment of methyl groups on nascent DNA after replication, primarily targeting hemimethylated substrates (10–12). DNMT3A/B performs *de novo* methylation of unmethylated, or hemimethylated substrates (13). DNMT3L, and some isoforms of DNMT3B lack the catalytic domains conserved in other DNMTs, but instead serves as an accessory protein, which stimulates methylation activity via DNMT3A (14, 15). Active demethylation occurs via Ten-Eleven Translocation (TET) proteins, which prime 5-mC (methylated cytosine) residues for removal via successive oxidation reactions(16). Decreased expression of TET proteins and low 5-hydroymethylcytosine (5-hmC) levels are associated with multiple cancer types (17) and developmental misregulation (18, 19). TET activity is associated with driving global and loci-specific demethylation in pluripotent cells. The activity of these systems of methyl “writers” and “erasers” compete with one another to regulate both benign and malignant phenotypes in cells (20).

Despite the relatively thorough understanding of the key enzymes controlling DNA methylation, open questions remain about how the global methylation landscape is established and maintained, in concert with other epigenetic and structural/genetic (sequence) mechanisms. Underlying genetic sequence context has been shown to play at least a significant contributing role in determining methylation, particularly local density of CpGs (21–25). Mathematical modeling has also shed light on the mechanistic bases for global bimodality in methylation, where so-called CpG collaboration, or interaction between multiple CpG substrates and the key enzymes, contributes to the nonlinear feedback promoting global bistability and epigenetic memory (25, 26).

In this work, we quantitatively analyzed the dependence of methylation on the presence of neighboring CpGs both in the context of annotated CGIs and genome-wide at single CpG resolution. We carried out the analysis in two human embryonic stem cell (hESC) cell lines in which genome-wide methylation profiling (WGBS) data is available, including cells in which key enzymes of the DNA methylation machinery have been knocked out (13, 27). These knockouts represent epigenetically perturbed systems, which lack methylating and/or demethylating power. We quantify the steep decrease of methylation with increased number or density of CpG neighbors in terms of two types of models: A Hill-function model supports the concept of local CpG density as a controller of an ultrasensitive switch in methylation. We theorize that this switch arises, at least in part, from the so-called futile cycle of opposing enzymes (namely DNMTs versus TETs), in analogy to classical zero-order ultrasensitivity(28). We measured quantitative shifts in the steepness and threshold of the switch in response to enzymatic perturbations and drift in methylation observed across cell generations, supporting this view. We further develop a novel approximation for a stochastic mathematical model of methylation dynamics involving CpG collaboration, enabling efficient parameter fitting. All in all, our results support a view in which dynamic interplay of the opposing writer/eraser methylation enzymes, together with local CpG density, determine methylation levels.

## Results

### A CpG Island’s Methylation Depends on its Size/Length

All datasets analyzed show characteristically bimodal methylation of islands. This feature has been understood to confer repression (high methylation) or activation (low methylation) to promoters (29). Figure 1 shows CGI-level methylation for HUES64 and HUES8 hESCs, along with IMR90 terminally differentiated fetal lung fibroblasts. While all cell lines exhibit bimodal behavior, IMR90 notably has more intermediate methylation.

**Fig. 1.**
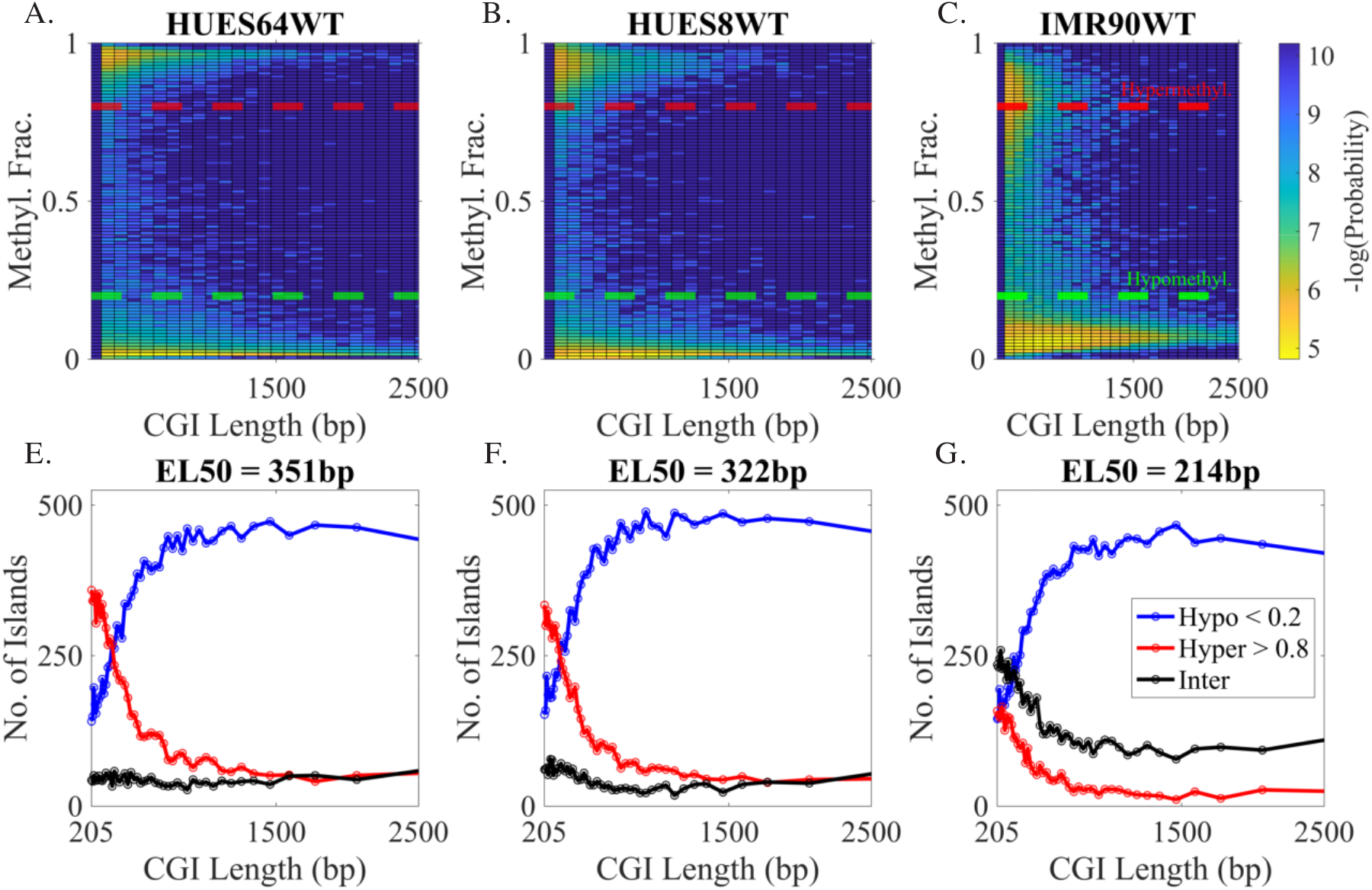
Dependence of CpG Island (CGI) methylation on CGI length in HUES64, HUES8, and IMR90 cells. *Top Row:* Bivariate heat maps of CGI-level averaging of bisulfite sequencing data comparing associated Methyl Fraction and Island Length; color corresponds to negative log-probability across binned tiles (yellow corresponds to highest density of islands). For each island, Methylation Fraction is defined as the sum over CpG-methylation values, divided by the number of CpGs in the island. CGI length is defined as the distance in bp between the first and last cytosine in CpGs associated to the island. *Bottom Row:* Curves display the number of CGIs that are classified into *Hypomethylated* (<20% Methylation Fraction, Dotted Blue Threshold), *Hypermethylated* (>80% Methylation Fraction, Dotted Red Threshold), and Intermediate methylation states, as a function of length. Length bins are determined by percentiles, giving an approximately equal number of islands in each length-bin. For each cell type, EL50 gives the length in bp at which the red and blue curves cross; that is, the length at which the number of islands classified into Hypo- and Hypermethylated groups is equal.

Bivariate heatmaps of CGI-level methylation versus CGI length indicate a shift of probability density toward lower methylation for longer islands. To characterize this shift, we binned CGIs according to length and then classified each CGI into *Hypomethylated* (<20% methylation fraction, dotted blue threshold), *Hypermethylated* (>80% methylation fraction, dotted red threshold), and Intermediate methylation levels. In all datasets, a clear trend was observed where the occurrence of hypomethylated CGIs increased with length (Figure 1, bottom). Very similar trends were observed when CGIs were classified according to their size (number of CpGs associated with the island) instead of their length, and CGI length and size were found to be linearly correlated (Fig S1-S2).

We defined the “Effective Length of CpG Island at 50% Probability of Hypomethylation”, EL50, as the length at which the blue and red curves cross; that is, the length at which an equal number of CGIs are classified as Hypo- versus Hyper-methylated. Similar EL50s were found in HUES64 (351 bp) and HUES8 (322 bp). Under this classification, the EL50 in IMR90 was shorter (214 bp, close to the minimum allowed size of CGIs). Notably, while the chosen thresholds (80/20) appear well-suited to HUES64 and HUES8, they do not fully capture the peaks (particularly the upper peak) in the bimodal landscape in IMR90. Nevertheless, IMR90 also shows an increased likelihood of hypomethylation of longer CGIs.

Even at longer CGI lengths, bimodality of CGI methylation in all cell types was evident, as a small population of CGIs persisted in the hypermethylated class even as the majority of islands at the longest lengthscales are hypomethylated. These results demonstrated that, while CGI methylation is bimodal at all lengthscales in the dataset, the relative populations in the two modes is linked to CGI length (or size). Increasing length of CGIs shifted the population from hypermethylated to hypomethylated.

### Discrete CpG Methylation Depends on Local Density, Recapitulating Trends at the CGI-Level

Classification of a region as an annotated CGI is stringent, in order to reasonably identify regions of particularly high CpG content. To test if the findings apply outside of CGIs, we classified all individual CpGs into hyper- or hypomethylation categories and analyzed them as a function of their local CpG density within a +/-50bp window.

An individual CpG was classified as hypermethylated if greater than 80% of reads at that CpG were methylated (and so on). CpGs were binned according to their discrete local density value (see Methods). The bimodality shown at the CGI-level and ensuing dependence of methylation were also observed at the resolution of local CpG density (Figure 2). Also similar to the CGI-level analysis, IMR90 had higher representation of Intermediate methylation.

**Fig. 2.**
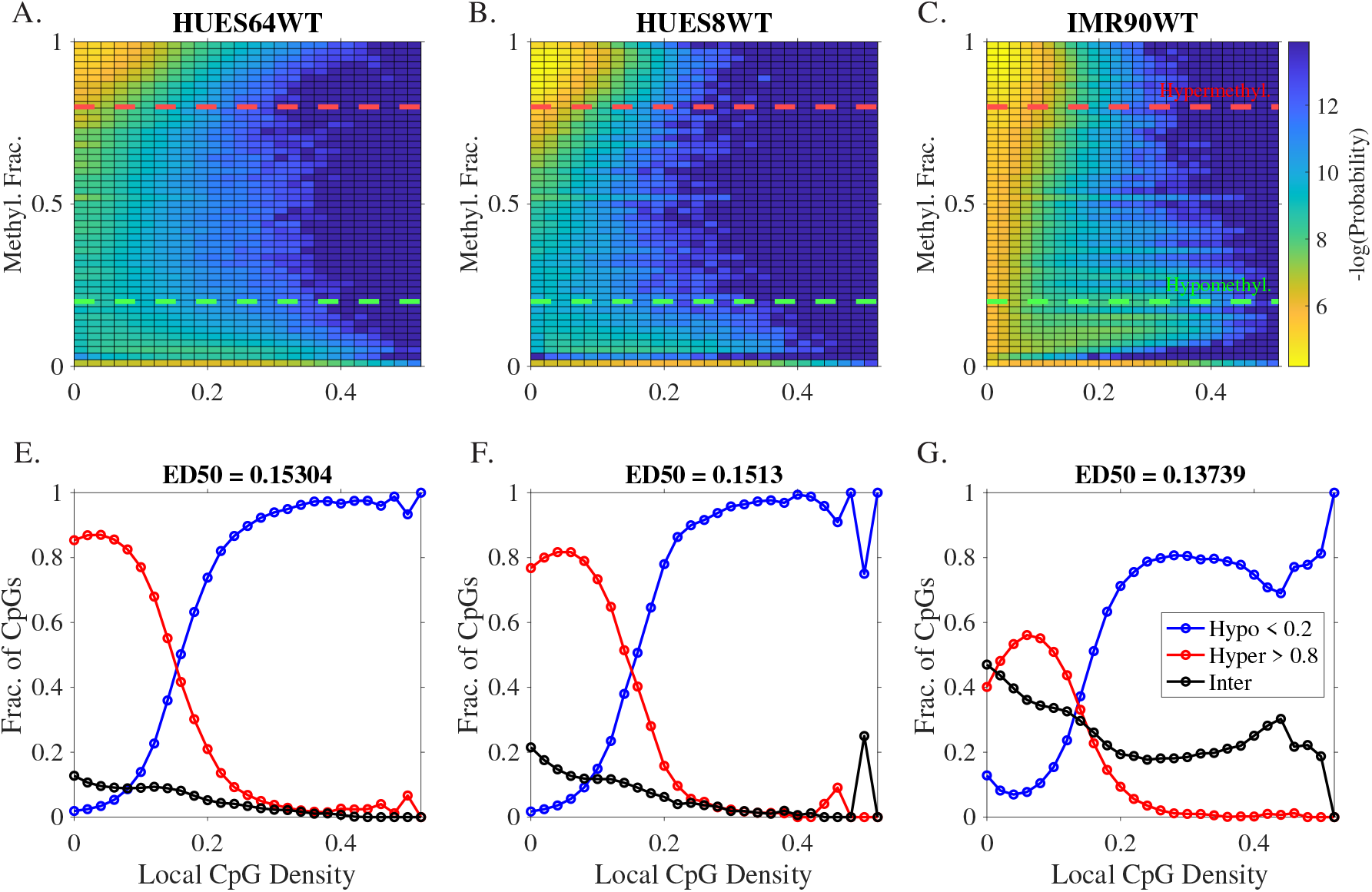
Dependendence of individual CpG-level methylation on normalized local CpG density with respect to a window size of 50bp (+/-). *Top Row* : Bivariate heat maps of individual CpG-level methylation from bisulfite data versus normalized local CpG density; color corresponds to negative log-probability across binned tiles (yellow corresponds to highest density of CpGs). *Bottom Row* : Curves display the fraction of CpGs that are classified into *Hypomethylated* (<20% Methyl Fraction, i.e. <20% of reads at that CpG are methylated, dotted green threshold), *Hypermethylated* (>80% Methyl Fraction, dotted red threshold), and Intermediate methylation groups, as a function of density. Each CpG in the dataset is associated to a discrete density value, without binning, based on the number of neighbor CpGs within a +/-50bp window. For each cell type, ED50 gives the density at which the red and blue curves cross; that is the density at which the fraction of CpGs classified into Hypo- and Hypermethylated groups is equal.

Individual CpGs with low local density (i.e., fewer neighboring CpGs in +/-50 bp) were more likely to be hypermethylated, whereas high-density CpGs were predominantly hypomethylated. In HUES64 and HUES8, the increase of hypomethylation with density was generally smooth and monotonic. Due to the higher prevalence of intermediate methylation, IMR90 presents a more complicated dependency on density, but the general trend of increased hypomethylation with density was the same.

Analagous to the EL50, we defined the “Expected Density of 50%” methylation, ED50, as the density at which an individual CpG is equally likely to be hypo- or hypermethylated. (When the crossover occurred between discrete density values, interpolation was performed). Similar ED50s were found for HUES64 and HUES8 (0.1530 and 0.15130, respectively). These correspond to an average of ≈7.6 neighbor CpGs in the window +/-50 bp. A somewhat shorter ED50 of 0.14 was found for IMR90.

In all, these results reveal that, at the level of individual CpGs regardless of genomic context, local CpG density is a strong predictor of methylation. Unlike in CGIs, where bimodality is evident at all lengths, bimodality of individual-CpG methylation was not evident at the highest density values, as few high-density CpGs are hypermethylated. At intermediate densities in all cell types studied, there was significant population of CpGs in both the hypo- and hypermethylated modes. (In IMR90, even at the lowest densities there was significant population in all three classes). Density values near the ED50 corresponded to the most pronounced bimodality of individual-CpG methylation.

### Effective Length and Density at 50/50 Probable Hypomethylation Point Shifts with Enzymatic Knockouts, Quantifying DNMT/TET Competition

We reanalyzed WGBS data from HUES64 cells harboring single- or double DNMT3A/B knockouts and from HUES8 cells harboring combinatorial DNMT/TET knockouts to ascertain how CGI-length- or CpG-density-dependent characteristics depend on the key enzymes controlling methylation dynamics. We repeated the classification of CGIs or CpGs and calculated the EL50 and ED50, as described above, in each cell type. We furthermore compared length/density curves as a function of passage number.

All of the analyzed knockout cells show shifts in the CGI Methylation versus CGI Length curves and in the CpG Methylation versus CpG-Density curves (Figures 3A-D). Double knockouts of DNMT3A/B (DKO) display increased representation of hypomethylated CGIs at all annotated CGI lengths, with this fraction of islands also increasing with time (i.e. number of passages) compared to the wild-type fraction (Figures 3A and 3C). At the individual CpG level, this behavior is seen predominantly at lower normalized CpG densities. Very similar shifts from WT are observed in HUES8 DKO as in HUES64 DKO cells. Conversely, HUES8 cells harboring triple knockouts of TET1-3 active demethylases (TKO) show decrease in hypomethylated CGIs at all lengths (along with an increase in hypermethylated CGIs).

**Fig. 3.**
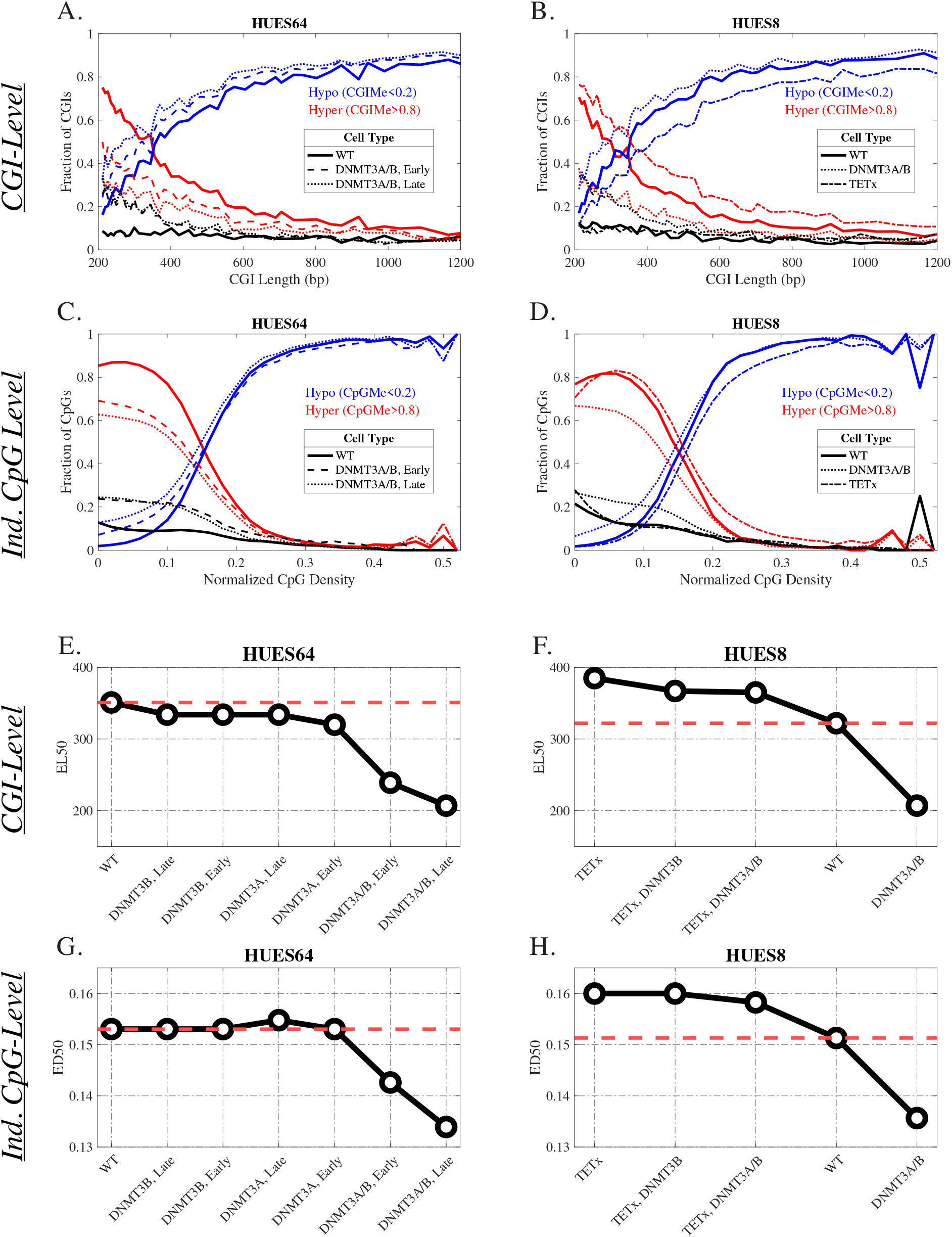
Comparison of CGI-Level (length) and Normalized Individual CpG-level (density) dependent methylation in HUES64 and HUES8 cells harboring DNMT3A/B Double Knockouts (DNMT3-DKO) or TET1-3 Triple Knockouts (TET-TKO). Wildtype (WT) HUES64 [A,C] and HUES8 [B,D] curves are the same as those in Figure 1 and Figure 2, with CGIs or CpGs classified into Hypomethylated (<20% Methylation, Blue), Hypermethylated (>80% Methylation, Red), and Intermediate methylation classes (Black). Overlain are corresponding curves from enzymatic knockouts. The legend gives the linestyle associated to the enzymes knocked out (e.g., TETx corresponds to TET-TKO cells). [E-H] Quantifying EL50 and ED50 in single- and double DNMT3A/B knockout systems in HUES64 cells and HUES8 DNMT/TET combinatorial knockout systems. Red dashed line denotes crossover values of the WT system for comparison. HUES64 DNMT3-DKO cells display marked decrease in both EL50 and ED50 compared to WT, and the difference increases with passage time (e.g. Early vs. Late, 15 cellular passages). DNMT3A or DNMT3B single-knockout systems do not display marked differences between one another, or with increasing passages in either EL/ED50 [E,G]. TETx knockout in HUES8 shows increased EL/ED50 compared to WT. HUES8 DNMT3-DKO shows decreased EL/ED50 and combinatorial TETx, DNMT3A/B knockouts show EL/ED50 values intermediate between WT and TET-TKO [F,H].

For both HUES8 and HUES64, DKO cells show a markedly decreased EL50 and ED50 relative to WT. In HUES64, EL/ED50 further decreases from Early to Late passage numbers. EL50s from single knockouts of DNMT3A and DNMT3B are only slightly decreased compared to WT, with no clear effect of passage number. ED50s for single knockouts in HUES64 show no or minimal change from WT. These results echo previous findings of minimal change of the methylation landscape in response to single DNMT3 knockout, whereas double knockout clearly decreases methylation (13).

In HUES8 cells, DKO cells show a similar marked decrease of both EL50 and ED50 compared to WT. TET1-3 TKO also shows a significant change but in the opposite direction, with EL50 and ED50 increasing compared to WT. Combinatorial knockout of TET1-3 with DNMT3B alone or with both DNMT3A/B knocked out show the calculated metric values shift back toward WT; similar trends are observed in ED50s. In all, these results show that the trends of CGI methylation versus CGI length, or CpG methylation versus CpG density, are broadly maintained under various perturbations to the methylation enzyme system, but are quantitatively shifted. The similar shifts in EL/ED50, toward higher or lower values in TET1-3 or DNMT3A/B knockouts, respectively, suggest that the EL/ED50 quantifies the point of balanced competition between the eraser and writer enzymes. Weakening the methyl “erasers” (TETs) enables CGIs to reach larger size/lengths (and similarly, CpGs to reach higher local densities), before becoming predominanty hypomethylated. Conversely, weakening the “writers” (DNMT3A/B, in this case) lead to predominant hypomethylation at shorter CGI lengths (and similarly, lower densities).

### Shorter CGIs Exhibit Time-Dependent Loss of Methylation Stability in DNMT3A/B-Double Knockouts

Further analyzing the effect of (de-)methylase knockout systems, unique CGIs were classified based on their change in methylation between wildtype and knockout. For each CGI captured in both cell types, in Figure 4, Δ *Methylation* is the difference in methylation fraction of the CGI in the first cell type, minus that in the second cell type. Each CGI is then classified in terms of its length and its methylation change. In all analyzed DKO systems, there was an enrichment of islands that experience moderate/drastic methylation loss at shorter CGI lengths (see also Fig. S6-S7). There was accumulating time-dependent loss of methylation, seen via differences between *Early* and *Late* passage timepoints (Fig. S5). The opposite effect, i.e., a CGI-size-dependent gain in methylation in the TET knockout system, was not clearly evident, supporting the notion that TKO-induced-methylation is broadly homogeneous at any size of methylated CGIs. However, further normalization to starting state (i.e., separating the analysis of CGIs in initial hyper/inter/hypo groups) shows that shorter, initially hypomethylated CGIs are more susceptible to TKO-induced increases in methylation, compared to longer CGIs (Fig. S7). Together, these results show that CGI size is a factor determining susceptibility to enzyme-knockout-induced methylation changes, particularly for methylation loss induced by DNMT3A/B knockout. These results also further support the idea that the CGI-length dependent trends of Figures 1-3 are dynamically maintained by enzyme competition, and that collectively, many CpGs in longer CGIs have more protected methylation status, whether hyper- or hypo-methylated, against fluctuations in enzyme levels.

**Fig. 4.**
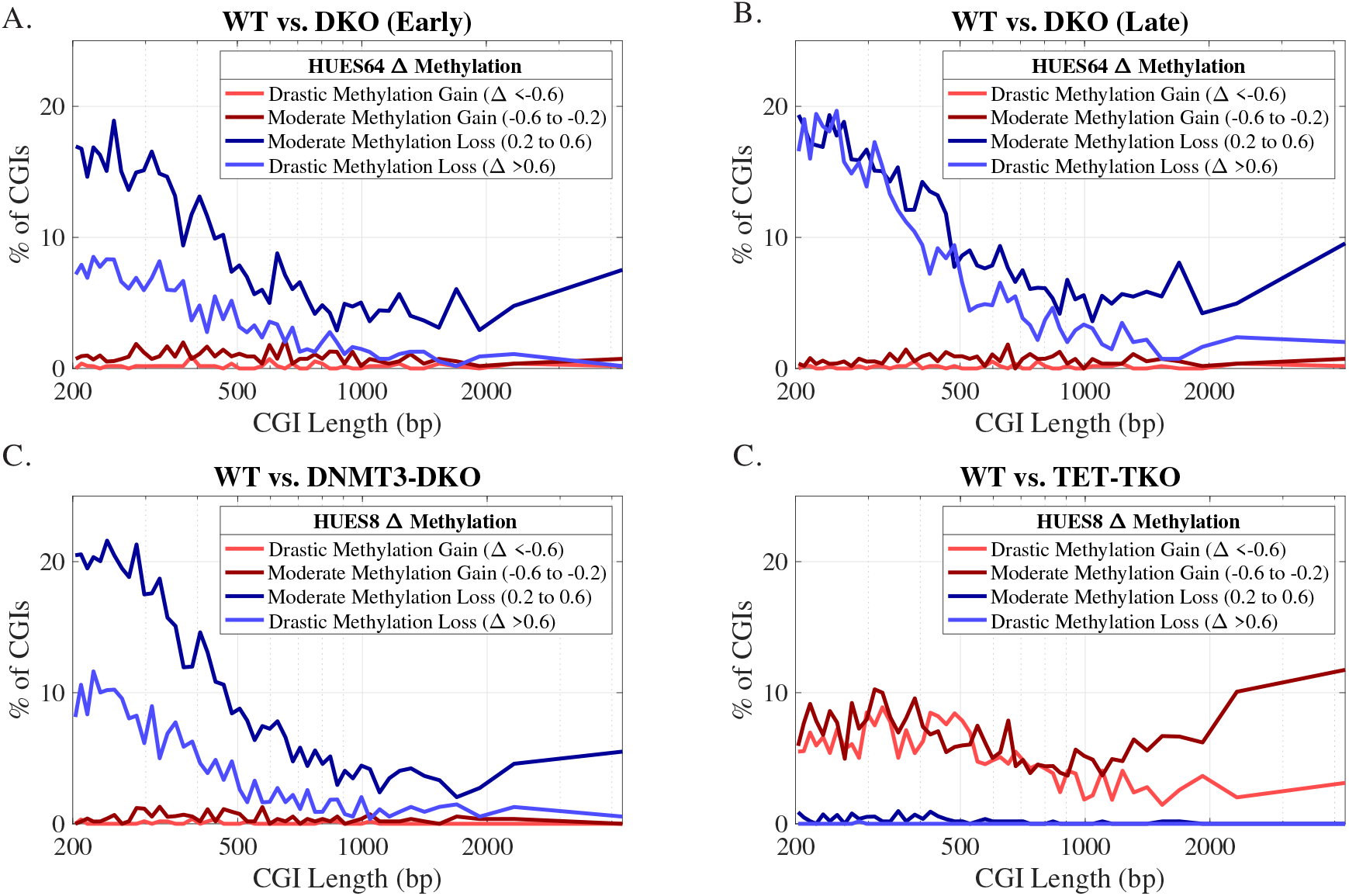
*Top Row:* Quantification of State-Switching CGIs in HUES64 DNMT3-DKO system. Panels calculate percentage of CGIs which experienced marked changes given the annotated cell types; islands which experienced less than a 20% change in methylation are omitted. In all shown comparisons, drastic and moderate methylation losses are associated with shorter CGIs, shown via decrease of these curves with increasing CGI length. (Early vs. Late: 15 cellular passages.) *Bottom Row:* Quantification of State-Switching CGIs in HUES8 DNMT3-DKO (*Left*) and TET-TKO (*Right*) systems. DKO system capitulates same susceptibility to methylation loss of shorter islands as shown in the HUES64 systems, but this size-dependence is not seen in the moderate/drastic gains of methylation exhibited in loss of de-methyltransferase activity in TET-TKO system.

### Fitting of Hill Function Parameters Exhibits Ultrasensitivity in CpG Methylation versus Local Density

Ultrasensitivity refers to switch-like behavior seen in a variety of biomolecular systems when a system exhibits a large change in output response on crossing some threshold of input (30, 31). Such a sigmoidal relationship between the input/output may be fit via a classical Hill function:

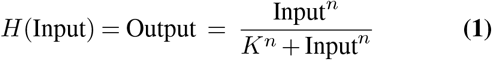

*Input* often refers to concentration of substrate. *K* is related to an equilibrium constant in idealized systems, but is more generally taken to be the *EC50*, or “Effective Concentration at 50% saturation”. We expanded upon this understanding through our analogous ED50 calculated metrics, using local CpG density in place of system concentration. Classical Michaelian kinetics has *n* (the so-called Hill coefficient) equal to 1. Ultrasensitve systems are characerized by *n>* 1 (32).

Exploring ultrasensitivity of CpG methylation with respect to local density, Figure 5 displays median, mean, 25-, and 75-percentile methylation curves for individual CpGs as a function of local CpG density in the aforementioned enzymatic knockout systems. These are the same data as plotted in Figures 3C-D, but do not rely on previously defined classification dependent on chosen cutoffs. We hypothesized that removal of any methyl writers or erasers would have a discernible effect on the system’s sensitivity to CpG density.

**Fig. 5.**
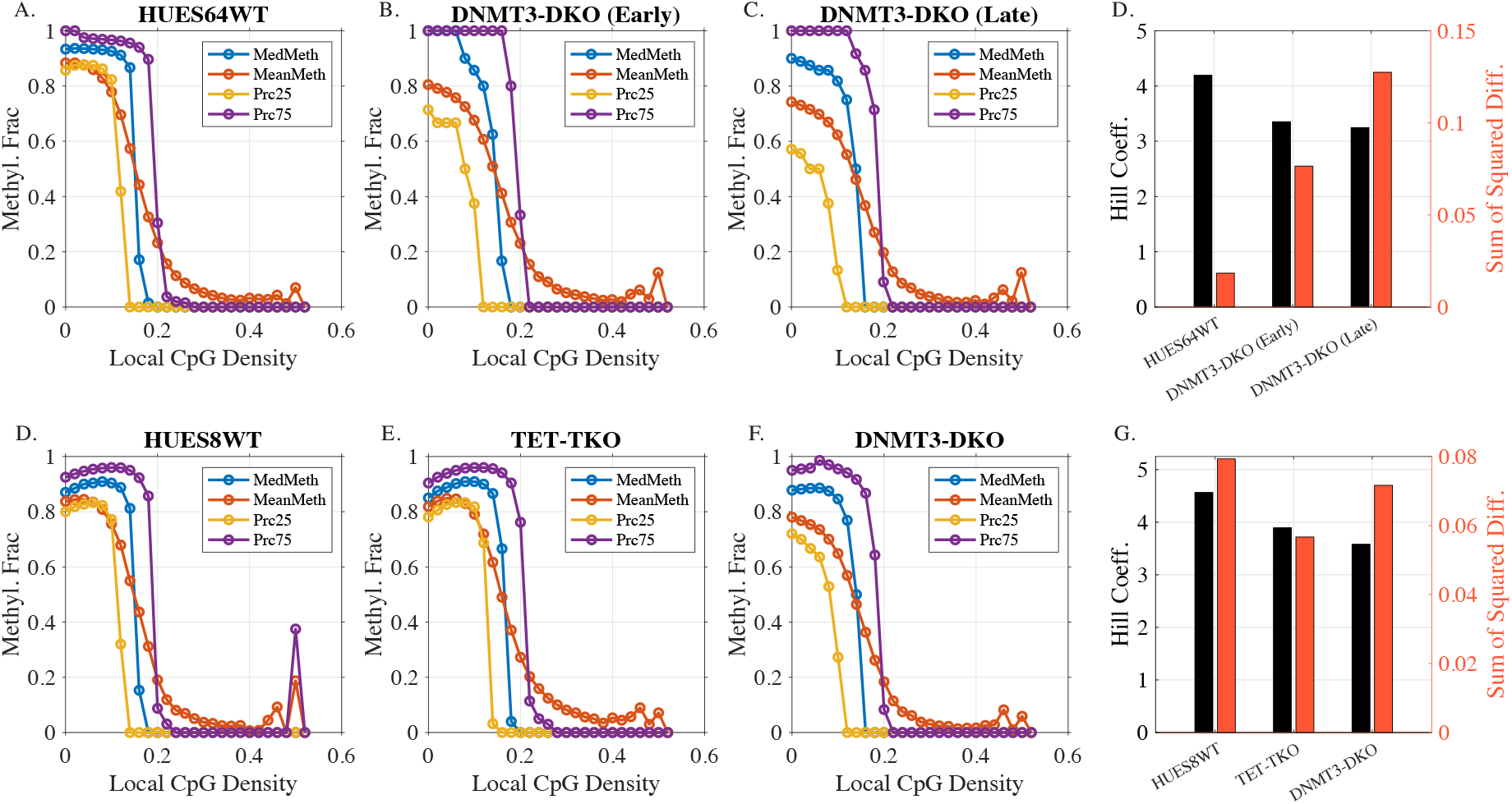
(*Top Row* = HUES64, *Bottom Row* = HUES8), (*Col. 1-3*): Individual CpG mean, median, 25th- and 75th-percentiles graphed as a function of normalized CpG density calculated within a +/-50bp window. (*Right Col*.) Mean Methylation Fraction curves (orange) are fit using a general Hill Function, with Hill Coefficients (i.e. exponents) and Sum of Squared Differences reported. Values of coefficients greater than 1 suggest the system exhibits ultrasensitivity with respect to CpG content. Fitted Hill coefficients of DNMT3A/B double knockouts, and TET1-3 triple knockouts methylation states decrease compared to the Wild-type.

In Figure 5, bimodality is apparent in that the median switches much more sharply than the mean, reflecting that population is shifting from one mode to the other. Remarkably, taking HUES64WT as an example, median CpG methylation decreases from 0.87 to 0.17, over a corresponding density change from 0.14 to 0.16. According to our density definition, this reflects the addition of a single additional CpG neighbor (from 7 neighbors to 8) within +/-50 bp. (Note that this switch occurs at density values in line with the ED50 values obtained from hypo/hyper curve crossing.)

Mean methylation curves (which also appear switch-like, though less steep than the median curves) are fitted to the Hill equation shown in Equation 1, reporting fitted coefficients and sum of squared differences. As mean methylation decreases with density, we fit to the function 1 − *H*(Input). An alternative method for estimating *n* was also employed; fitted model curves for both methods overlain with mean methylation, along with parameterization are shown in Fig. S8-S11. The resulting Hill coefficients illustrate that WT HUES64/8 cells are therefore ultrasensitive with respect to CpG density, with a high Hill coefficient *>* 4 (Median methylation curves show even greater ultrasensitivity, with Hill coefficients ranging between ≈ 8 − 30). We also fit *n* for somatic IMR90 cells; while IMR90 showed a more complex density-dependent methylation, the significant, steep decrease of mean methylation that occurs near a density value of 0.14 is fit by *n* = 4.1 (Fig. S11), similar to the value in the hESCs.

In the hESCs, when subject to either DKO or TKO modulation, there is a decrease in the ultrasensitivity to CpG substrate density compared to the WT, as evidenced by lower fitted *n*. This supports the hypothesis that removal of CpG methyl writers and/or erasers reduces the cell’s ability to maintain ultrasensitive, switch-like behavior required for regulated bimodal gene expression. Together, these results unveil the idea that CpG density functions as an ultrasensitive methylation switch.

### Mathematical Models Including Collaborative Mechanisms Better Discern Experimentally-Observed Density-Dependent Trends in Methylation

We performed mathematical modeling to investigate the mechanistic basis for the observed ultrasensitive methylation switch, and for the enzyme-knockout-induced shifts in these trends. Our model is based on a previous framework employing CpG *collaboration*, which refers to the effect of nearby CpGs on rates of enzymatic reactions at a given CpG (26). The model is shown schematically in Figure 6A and described in Methods. Briefly, the model assumes that each CpG exists in one of unmethylated (u), hemimethylated (h), or methylated (m) states. Transitions between these states occur either independently or collaboratively (dependent on the presence and methylation status of neighbor CpGs), with specified kinetic constants. Demethylating reactions also occur passively due to DNA replication, dependent on the length of the mitotic cell cycle. The “Standard” model includes only CpG-independent transitions (including replication-associated methylation loss). The “Collaborative” model adds the distance-dependent neighbor effects on transition rates. The full, stochastic model requires simulation of many trajectories to achieve good estimates of model-predicted methylation distributions, and this poses a challenge for parameter fitting against data. To address this, we developed a coarse-grained approximation based on collective CpG “macrostates”. The steady-state distribution is then directly computed using the framework of the Chemical Master Equation (33, 34).

**Fig. 6.**
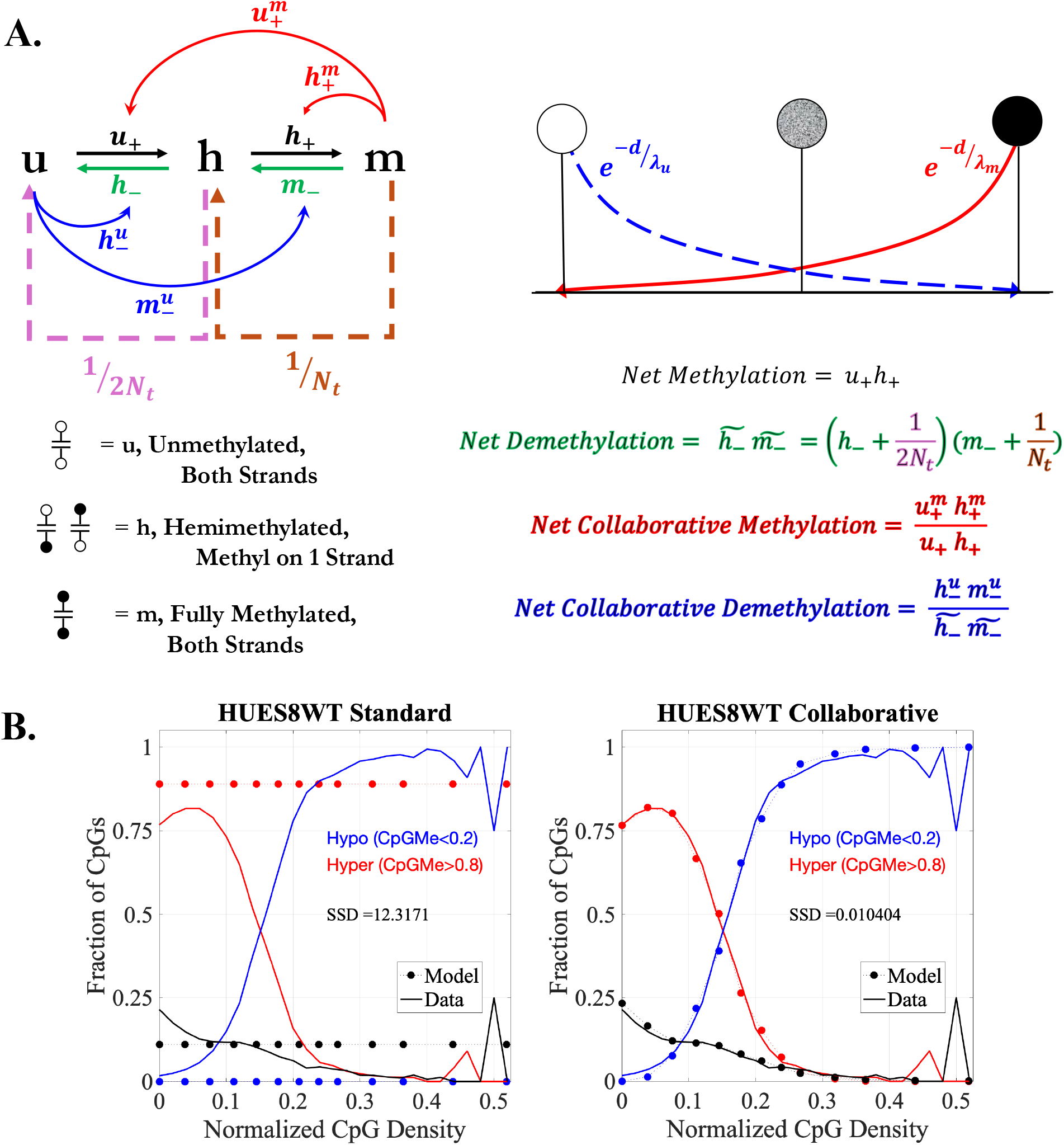
*A:* Model Schematic of DNA Methyl State Transitions and Consideration of Exponential Inter-CpG Distance Dependence. *B:* HUES8 Methylation data overlain with Mathematical Model as a function of local CpG Density. Data and Model outputs are classified in to Hypo/Hypermethylated bins as described previously, and Sum of Squared Differences is reported as a metric of Goodness-of-Fit.

HUES8 WT methylation data for individual CpGs are fit using the Standard and Collaborative Mathematical Models (Figure 6B). For a given set of model parameters in the fitting, the methylation distribution is computed for a group of CpGs, varying inter-CpG spacing to simulate changing local density. The Standard model, with no consideration for collaborating/interacting methylation events, is unable to exhibit any density dependence, since every CpG in the model independently reaches the same steady state probabilities (*p*_*u*_, *p*_*h*_, *p*_*m*_), regardless of neighbor context.

In contrast, the Collaborative model, employing an exponentially decreasing dependence on inter-CpG distance to simulate locally cooperative enzymatic activity, achieves an excellent fit to the experimental data curves. These results support that CpG collaboration is responsible for the observed methylation switch. The results further support that this modeling framework, which does not directly treat any details of genomic context or other layers of the epigenome (histone marks, chromatin structure, etc.) other than CpG sequence context, is sufficient to fit experimentally observed trends.

### Parameterization of DNMT/TET-KO Methylation Landscapes Provides Evidence for Tunable Ultrasensitivity via Rates and Lengthscales of Enzymatic Action

Using the Collaborative model, fitting was performed on the HUES8 DKO and TKO cell systems to understand how the shifted curves in the enzyme knockouts could relate to molecular parameters. Parameter sets giving good fits, quantified via the reported sum-of-squared distances, were obtained for all three HUES8 cell types (Fig.7A). However, parameter sets were not unique. To compare different parameter sets, three collective measures were used. Net (de-)methylation refers to the net flux of (de-)methylating reactions in the Standard model. Net collaborative (de-)methylation refers to the net flux of collaborative (de-)methylating reactions, scaled by the corresponding non-collaborative net flux in the same direction. As such, this gives a measure of the maximum relative amount of methylating (or demethylating) activity that is collaborative in nature. Because of the distance-dependence of collaborative reactions, this contribution decreases with decreasing distance of neighbors. Finally, the lengthscale of (de-)methylation refers to parameters λ_*m*_ and λ_*u*_, which are the decay constants for the exponential distance functions.

**Fig. 7.**
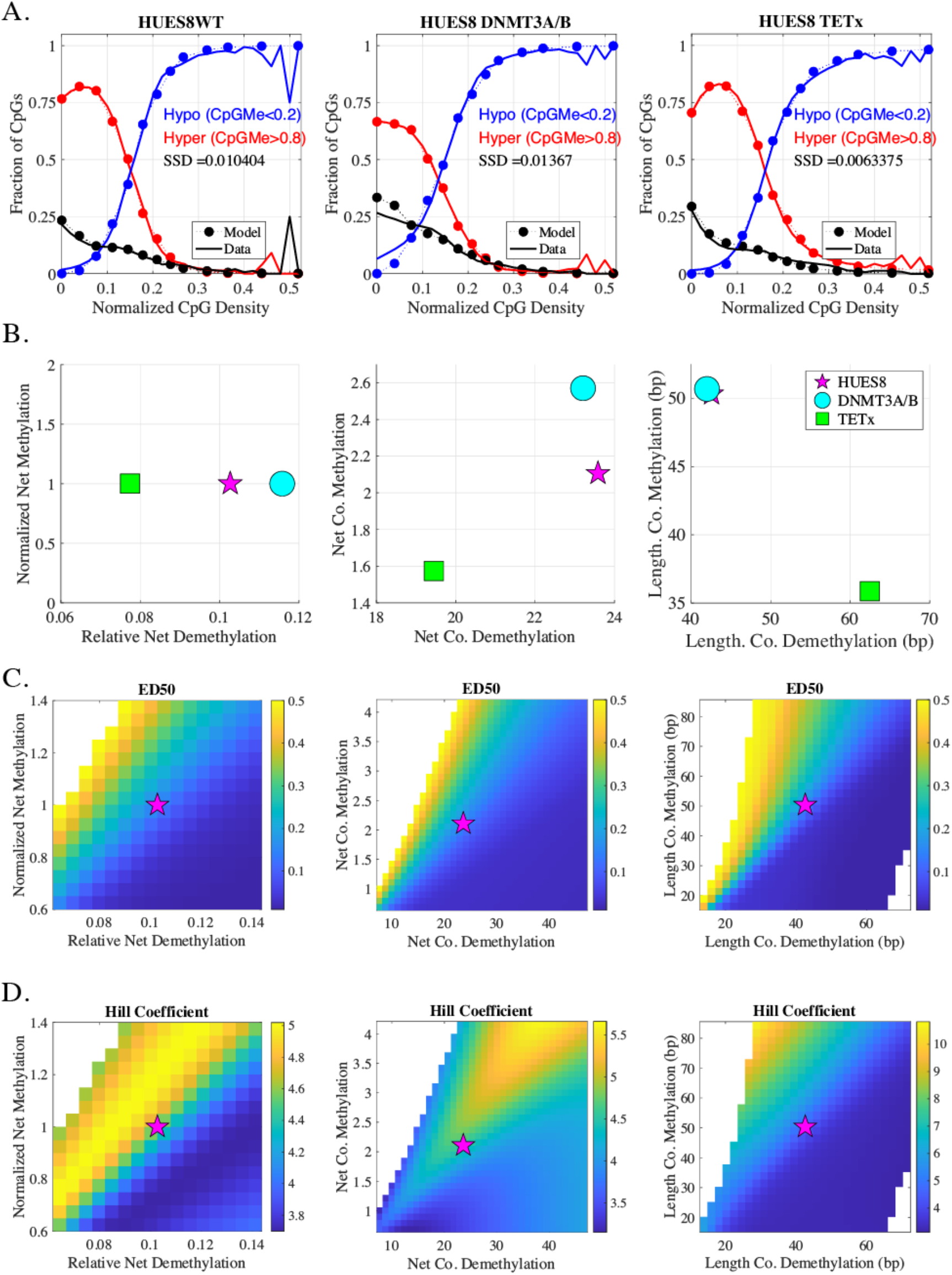
*A:* Representative mathematical model fits to three HUES8 variants (WT and enzyme knockouts). *B:* Comparison of fitted model parameters for the three cell variants. Parameter changes are summarized in three planes, using three general metrics: Net (De-)Methylation, Net Co. (De-)Methylation, and Lengthscale of (De-)Methylation. See Methods and Figure 6 for detailed definitions. (*C-D:*) A wider range of parameter changes are shown, and their effect on the density curves. Each plane shows the effect of varying parameters on either ED50 (*C*), or steepness (Hill coefficient, *D*). For each column, parameters are varied in-plane, while all other parameters are held fixed. Magenta star represents best fit to HUES8WT, and parameter sweeps are centered on this point in parameter space (see Methods).

For the representative fits shown in Fig.7A and B, the TETx knockout shows decreased net demethylation rate (measured relative to net methylation rate), as expected. Similarly, the DNMT3A/B knockout shows increased relative net demethylation rate, as expected. However, good fits were not achieved by varying these net forward/reverse reaction rates alone. Parameter fitting suggests that the knockout lines also have modulated collaborative enzyme parameters. The TETx knockout showed significantly decreased rates of both methylating and demethylating collaborative reactions, while the

DNMT3A/B knockout showed a more moderate decrease in the collaborative methylation measure, and a minimal change (increase) in collaborative demethylation. Finally, the model fits suggest that lengthscales of collaborative reactions are unchanged in DNMT3A/B knockout, whereas they are significantly changed in the TETx knockout (increased λ_*u*_, decreased λ_*m*_).

Varying each collective parameter measure independently reveals effects on both steepness (Hill coefficient, *n*) and threshold (ED50) of the methylation switch (Fig.7C and D). Parameter sweeps were performed around the optimized HUES8WT values; keeping all else equal. For example, to vary the net methylation, given by *k*_*u*+_*k*_*h*+_, by a factor *s*, each of the individual parameters is scaled identically by 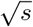. In all, the parameter sweeps reveal how all three collective measures tune the characteristics of the switch. In general, weakening methylation parameters along any measure decreases ED50, in line with the experimental observations for DNMT3A/B knockout. Conversely, weakening demethylation parameters increases ED50, in line with TETx knockout observations. The Hill coefficient shows more complex behavior with respect to the parameters, but two observations emerge. First, simultaneously increasing net collaborative rates (both methylating and demethylating) leads to even higher Hill coefficients than were observed from experiments. This is reminiscent of the classic bistable genetic switch (35), in that increasing positive feedback on the two modes increases the nonlinearity of the switch (here, the positive feedback is imposed by the collaborative terms on both sides). Another observation is that the highest theoretical Hill coefficients (*>* 10) were obtained in a regime where the lengthscale λ_*m*_ was long (≈ 80 bp) while λ_*u*_ was moderate (≈ 30 bp). While further study is needed to understand how the model parameters relate to specific molecular mechanisms, these results suggest that the steepness and threshold for the methylation switch are controlled by the interplay of antagonistic enzymatic rates and lengthscales on which their activities are sensitive to the surrounding context of the central CpG.

## Discussion

### Creation of Flexible Bioinformatic/Modeling Tool to Infer Molecular Mechanisms

In this work, we combined quantitative analysis of DNA methylation dependence on local CpG context (either in CpG islands or genome wide) with mathematical modeling to characterize the dynamic interplay between writing and erasing enzymes in a given cell type. We found that the length- or density-dependent methylation trend can be characterized as an ultrasensitive switch. In hESCs, quantitative shifts in this switch occurred with knockouts of either DNMT3A/B or TET1-3. Enzymatic perturbation affected both the threshold (EL50 or ED50) and steepness (Hill coefficient) of the methylation curves relative to WT. Mathematical modeling suggests that the methylation switch indicates collaboration among CpGs, i.e., where the methylation status of nearby CpGs affects both methylating and demethylating reaction rates at a given CpG. The shift in ED50 (to higher/lower density in TET1-3 and DNMT3AB knockout, respectively) results from a shifted balance of opposing enzymes. Both TET1-3 and DNMT3A/B knockouts showed reduced Hill coefficients, indicating a general reduction of the collaborative strength. Our analyses show that behaviors at broad genomic features, e.g. CGIs, may connect to those shown at the individual CpG level and allow researchers to elucidate broader determinants of regulation by disentangling the primary drivers at each of these levels. In the future, this approach may be used not only for embryonic stem cells, but to also investigate modulation in DNA methylation due to differentiation, disease modalities such as cancer, aging, and effects of drug perturbations on the regulatory landscape.

### Ultrasensitivity of DNA Methylation with CpG Density Resulting from Opposing Enzymes

We found that DNA methylation exhibits an ultrasensitive switch-like dependence on CpG density, with a Hill coefficient *>* 4. DNA methylation machinery may be considered to participate in a “futile cycle”, since there are two separate, opposing enzyme-catalyzed reactions operating on the same substrate in opposite directions (i.e., broadly, the DNMT-catalyzed “writing”, versus the TET-catalyzed “erasing”). Futile cycles are widespread in biology, with classic examples being in metabolism (e.g., glycolysis versus gluconeogenesis) and signaling (phosphorylation/de-phosphorylation cascades)(36). The direction of net flux in a futile cycle is controlled by differential regulation of the opposing enzymes(37).

In their seminal work(28), Goldbeter and Koshloand proposed that this type of control could achieve extreme, switch-like ultrasensitivity (which they termed “zero-order ultrasensitivity” under conditions of enzyme saturation) wherein the net flux switches directions abruptly at a threshold ratio in concentrations of the opposing enzymes. This mechanism is thought to be one contributing factor underlying observed ultrasensitivity in various systems (38–41) (reviewed in (42)). We propose that the ultrasensitivity we observed in DNA methylation versus CpG density is another example of zero-order ultrasensitivity, although the impact of the sequence context of the CpG substrate on kinetics also makes it distinct from other systems. Our empirical ED50 reflects the density at which, under the original theory, *k*_1_*E*_1*T*_ = *k*_2_*E*_2*T*_, that is, the products of the effective rates and total enzyme concentrations are equal. Here, reaction 1 represents the net writing reactions, 2 represents net erasing reactions (or vice versa), and under our model, the switch is achieved not by changes in the enzyme concentrations per se, but by the dependence of the effective net rates (*k*_1_ and *k*_2_) on CpG density. Though our full models are more complex and involve many more parameters, we confirmed that the model-predicted ED50 fulfills this equality. A prediction of the theory is that changes in relevant enzyme concentrations should shift the ED50; for example, decreasing TETs should increase ED50, since a higher CpG density is required for the net fluxes to equalize. Indeed, this prediction is confirmed by our results for the DNMT3A/B and TET1-3 knockout cell lines, which show the predicted directional shifts (Figures 3 B,D). These results further support the conclusion that the ultrasensitive switch in DNA methylation is explainable by an adapted form of the Goldbeter-Koshland theory, pointing to a mechanism whereby the interplay of CpG density and differential expression of writing/erasing enzymes can achieve relatively targeted methylation modifications. This view is further supported by our finding that susceptibility of CGIs to methylation change as a result of enzyme perturbations is not uniform, but rather dependent on their relative size/length (Figure 4). In all, our results suggest that the methylation futile cycle, combined with the enzymes’ sensitivities to local CpG sequence context, is the major driver of globally bimodal methylation landscapes.

### Relationship Between CpG Density and DNA Methylation

The relationship between local CpG density and DNA methylation has been studied previously, with most reports supporting a general inverse relationship. CpG Islands (CGIs), which have relatively high local density of CpGs, are recognized to be predominantly hypomethylated(43). An early genome-scale study pointed to increased methylation levels of CpGs with decreased local CpG density(44). Within CGIs specifically, CpG-rich (as well as GC-rich) sequence patterns were found to be over-represented in unmethylated (versus methylated) islands(45, 46). Relatedly, under a genome-scale definition of “CpG clusters” in human DNA, Lovkvist et al. reported that clusters with fewer or less densely spaced CpGs are predominantly hypermethylated, while larger clusters are hypomethylated (25). In a variety of human and mouse tissues, when CpGs were binned by density, genome-wide, in a 2500 bp sliding window, a strong negative correlation was found between CpG density and methylation(21). In Partially Methylated Domains, flanking sequence context, and especially CpG content near an individual CpG, was shown to be predictive of methylation, although one study reported increased methylation with increased local (+/-20 bp) CpG density(22), while another reported the opposite trend (decreased methylation with increased local (+/-35 bp) CpG density)(23). An earlier study reported that methylation initially increases with increasing CpG density, but then sharply decreases again at higher densities.(24)

Our approach to characterizing the genome-wide relationship between local CpG density and CpG methylation, regardless of other genomic context, by CpG binning is similar to that of Chen, et al. (21), although we focus on individual CpGs (rather than sliding-window averages) and a smaller local neighborhood (+/-50 bp here). Our mean methylation curves (in Figure 5) appear qualitatively very similar to theirs, and they report similar patterns across a number of human and mouse tissues. Another study characterized the decrease of methylation with CpG density in mouse DNA, finding a half-maximum point near 8 CpGs per 100 bp, which is very close to the ED50s we report here(47). In our results and theirs, there is a slight initial increase of methylation at low densities, followed by a sharp drop at higher densities, reminiscent of the finding of Edwards et al. (24). In our study, this feature is somewhat cell-type dependent and is particularly apparent in IMR90 cells. Our modeling indicates that the initial increase is due to collaborative methylation, which dominates at lower densities because the non-collaborative reactions also favor methylation. These results suggest that a non-monotonic relationship between CpG density and methylation is present when density is quantified in a fine-grained manner, which may also help explain the seemingly contradictory findings of neighbor-CpG influence in PMDs(22, 23). However, in our study all cell types analyzed shared a similar ultrasensitive switch (drop) in methylation at higher densities, broadly in agreement with the general trends uncovered in previous genome-wide or genome-scale analyses (21, 25, 44).

### Molecular Mechanisms of Collaboration

“Collaboration” in this work, adopted from Haerter, et al. (26), refers to any mechanism by which CpGs (either directly or indirectly) influence the rates of enzymatic writing/erasing at another CpG, possibly in a manner that depends on their methylation state. Various molecular mechanisms may underlie this collaboration. For example, DNMT1 is preferentially recruited to sites with more neighboring hemi-methylated sites (48), and these CpGs show faster methylation rates (49). The latter finding is consistent with DNMT1 processivity, where the enzyme can successively methylate neighboring sites. DNMT1 processivity has been characterized *in vitro* (50–55), and indirectly *in vivo*, evidenced by locally correlated maintenance methylation kinetics(56, 57). UHRF1 helps recruit DNMT1 to hemimethylated CpGs(58–60). Monoubiquinated histone H3 helps recruit DNMT1 to DNA stretches with multiple, but not one, hemimethylated CpGs (48), supporting the concept that UHRF1 may preferentially direct DNMT1 to CpG-dense regions.

Active demethylation processes carried out by TET enzymes also show some evidence of collaboration. DNA strand sliding by a TET protein was observed by high-speed Atomic Force Microscopy (AFM) in a DNA origami nanochip(61), though another study saw no evidence of this type of processivity (62). The binding domains of TET1 and TET3 prefer CpG-rich regions (63, 64) and TET1 binding correlates with CpG density (63, 65, 66).

### Insights from Mathematical Modeling

The mathematical model gives insight to the mechanistic basis for the genome-wide CpG-density-dependent methylation curves. Broadly, the model suggests that antagonistic methylation/de-methylation processes occur continuously at every CpG, with the CpG-density-context of each CpG dictating the relative rates of these opposing processes. However, density-dependent rates are alone insufficient to give bimodality; rather, interactions among neighboring CpGs provide the necessary positive feedback to stabilize either hypo- or hypermethylated states. Previous studies of the competition between TETs and DNMTs have also pointed to enzymatic competition as a central determinant of steady-state methylation levels, and have uncovered different relative activity levels in different genomic contexts using mathematical models, such as in high/low or unmethylated regions (67), or in, e.g., enhancers or Transcription Factor Binding sites (68). In the present work, our model analyzes enzymatic activities solely based on sequence-encoded CpG positions. Future studies are needed to disentangle the relative roles of genetic sequence versus other local context information (histone marks, relative positioning compared to enhancers, promoters, gene bodies, etc.) in determining enzymatic recruitment and local activities as well as ensuing methylation patterns. Our models also cannot distinguish between direct or indirect influence of local sequence on the enzymatic behavior.

Our model was adapted from previous works(25, 26) and modified in key ways. First, we have formulated the model as a stochastic chemical reaction network, with kinetic rates as free parameters. Second, we devised a mean field approximation to enable rapid solution of the steady-state distribution over methylation states. This approach enabled us to efficiently calculate the methylation distribution as a function of neighbor-CpG-density without simulating many trajectories, enabling investigation of a broad parameter space for relatively large system sizes. Third, we used the genome-wide, cell-type-specific distribution of CpG-methylation classes (hypo/hyper/inter) as a function of local-CpG-density to fit the model parameters. We also investigated how the key parameter metrics (net (de)-methylation rate, relative collaborative (de-)methylation, and lengthscale of collaborative (de)-methylation) control the threshold and steepness of the methylation switch. We found that purely exponential distance functions (i.e., decreasing over a lengthscale determined by λ) were sufficient to recapitulate the experiment-derived density curves, while Lovkvist, et al.(25) suggested that collaborative methylation reactions have a more long-ranged, power-law dependence on distance.

### Lengthscale of CpG Interactions

Although the parameters cannot be unambiguously determined, good model fits were achieved with collaboration lengthscales on the order of 40-65 bp for the exponential distance functions. These values are in line with our previous work supporting a processive lengthscale of DNMT1 of 38 bp in hESCs, based on exponential decay of kinetic correlation (57). However, some genomic contexts (especially CGIs and TFBRs) showed more pronounced deviation from exponential correlation than others, with evidence of more long-range effects. Other experimental evidence supports a role for relatively short-range interactions. For example, it was found that CpG density within windows of ±35bp was optimal for predicting PMD-specific hypomethylation (23). In a study of the effect of local sequence context on CpG methylation(22), the strongest determinant at a central CpG was the presence of adjacent CpGs, although other neighboring CpGs had an impact on methylation that decreased continuously with distance to roughly +/-40 bp. More work is required to further elucidate the lengthscales, distance-dependence, and mechanisms of CpG collaborative interactions.

It is important to note that our modeling, while providing good fits to the global methylation trend, do not take into account many details of local epigenetic/chromatin context.

One study found that, while CG density alone is a significant determinant of methylation genome wide, in CGIs there was a principal role for additional factors such as transcription factor binding sites (47). In the future, the analysis and modeling approach presented here could be augmented with more detailed genomic/epigenomic context factors to better predict methylation levels.

## Methods

### Cell Line Experimental Data and Sources

Thirteen Whole Genome Bisulfite Sequencing (WGBS) datasets from human cells were reanalyzed. All figures and analyses are based on either CpG Islands (CGIs) across the genome, or all captured CpGs from Chromosome 1. Cell lineages and GEO Accession numbers are listed in Table S1. Data are from human Embryonic Stem Cell lines HUES64 and HUES8 and a fetal lung fibroblast cell, IMR90. The hESC datasets include varying types and passage numbers of enzyme knockouts, including single knockouts of DNMT3A and DNMT3B along with DNMT3A/B double knockouts (DKO) in HUES64(13). Furthermore, DNMT3A/B double knockouts (DKO) and TET1-3 triple knockouts (TKO) were analyzed in HUES8 (27).

### Characterization of CpG Islands and Local CpG Density

Human genome assembly (*hg19*) and CpG Island annotations are acquired using the UCSC Genome Browser (69), where islands are considered if the ensuing genome segment meets the following criteria: *(i)* Guanine-Cytosine content greater than 50%, *(ii)* minimum segment length of 200bp, and *(iii)* the following inequality:

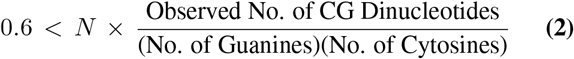

where *N* is the length (in base-pairs, bp) of the considered DNA segment. For a given CGI, CGI-Level Methylation Fraction is defined as the average over the methyl fractions (also termed methylation ratios) of the CpGs of which it is comprised. For classification of CGIs, methylation fraction above 0.8 is defined as Hypermethylated, below 0.2 is defined as Hypomethylated, and islands in between are denoted as Intermediate. CGI length (distance in bp between the cytosines of the first and last associated CpG) is used as a metric for general CGI size.

A normalized metric of localized CpG density, around a given CpG, is defined as:

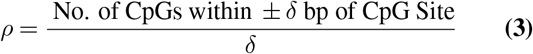

In this equation, *δ* represents the window size (+/-) around an individual CpG. As a normalization constant, we also use *δ*, reasoning that it represents the theoretical maximum number of CpG neighbors that could be possible given 100% CpG content. A value of *δ* = 50 bp is used.

### Calculated Threshold Metrics: *EL50* and *ED50*

The EL50 describes the lengthscale in bp at which it is equally probable to find a CGI in either the high- or low-methylation macrostates (i.e., hypermethylated versus hypomethylated, irrespective of the CGIs in the intermediate category). Similarly, the ED50 signifies the density value at which it is equally probable to find a single CpG in either the high- or low-methylation fractions. Explicitly, these metrics denote the crossover point of hypermethylated and hypomethylated annotated CGIs or fraction of classified CpGs (see Row 2 of Figures 1, 2, or Fig. S2). Note that a related but slightly different definition of EL/ED50 is by the CGI length (or CpG density) of half-maximal methylation fraction. The latter definition does not depend on threshold classification of hyper- versus hypomethylation.

### Mathematical Model Description and Fitting to Data

The model, adapted from Haerter et al. (26), is comprised of “standard” reactions, containing only linear transitions of CpG states, “collaborative” reactions, simulating non-linear transitions via distance-dependent bystander reactions, and cyclic replication events.

### Model Overview: Standard Model

The model assumes that each CpG can be in one of three states: unmethylated (*u*), hemi-methylated (*h*), or methylated (*m*). The basic reactions for gain and loss of methyl marks are thus:

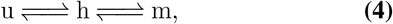

with separate rates for each reaction, denoted as {*k*_*u*+_, *k*_*h*−_, *k*_*h*+_, *k*_*m*−_}. (Subscripts denote the starting state of the site and the direction of the reaction, i.e., whether a methyl group is added (+) or removed (-)). The methylating reactions, u → h and h → m, are assumed to be catalyzed by methyltransferases, with DNA methyltransferases 3 A/B (DNMT3A and B) regarded as responsible for *de novo* methylation, and DNMT1 primarily responsible for maintenance methylation over mitotic cycles. DNMT1 shows a high preference for the hemi-methylated CpG substrate (70), and thus plays a central role in catalyzing h → m post-DNA-replication, but the roles of DNMT3A/B versus DNMT1 in *de novo* versus maintenance methylation cannot be clearly separated (71).

The demethylating reactions, m → h and h → u, are assumed to happen either actively, (catalyzed mainly by TET1-3 enzymes (72)) or passively, as a result of DNA replication, where cytosine on the nascent strand is unmethylated initially. The classical model of DNA methylation inheritance(73, 74) gives that every methylated CpG becomes hemi-methylated in both daughter cells (until DNMT1 acts to re-assert symmetric methylation), and every hemi-methylated CpG yields one un-methylated and one hemi-methylated daughter CpG. Mathematically, we adopt a “continuous replication” approximation(26), in which this passive methylation loss can be captured through rate constants linked to the cell cycle time, *Nt*, by:

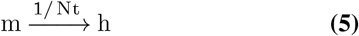

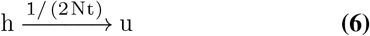

Incorporating this passive demethylation into the reaction scheme of Eq.4, the rate constants become 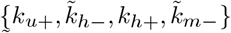, where the combined rate constants 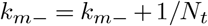 and 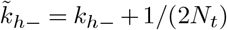 incorporate both active and passive demethylation.

### Model Overview: Collaborative Model

A number of mathematical modeling studies have provided support for the presence of interactions (also called “collaboration”) among CpGs in dynamic methylation processes, including in maintenance, *de novo* methylation, and demethylation reactions (26, 57, 75–77). We modeled a system of *N* CpGs as a non-uniform 1D lattice (i.e., the distances between neighbor CpGs need not be uniform). We assume that a CpG in a given state can impact the rate of a reaction on a nearby CpG via a distance function, *f* (*d*), where *d* is the inter-CpG distance in bp. We include the following collaborative reactions: {*u*+^*m*^, *h*+^*m*^, *h*− ^*u*^, *m*− ^*u*^}, where, for example, *u*+^*m*^ denotes a CpG in the *u* state that is methylated on one strand to become *h*, in a way that is dependent on a nearby CpG in the *m* state. Our model assumes that the collaborative methylating reactions ({*u*+^*m*^, *h*+^*m*^}) are governed by an exponential distance function, 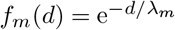. Likewise, the collaborative demethylating reactions (*h*− ^*u*^, *m*−^*u*^) have distance function 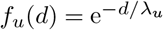. Adding in these collaborative reactions, the total rate for a given CpG at position *i* to undergo the u → h reaction becomes:

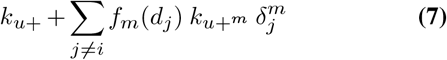

where *k*_*u*+_ is the independent-site (standard) rate constant, *k*_*u*+_*m* is the collaborative rate constant, and 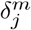 is an indicator that is 1 when the *j*th CpG is in the *m* state, and 0 otherwise. The *j*th CpG is distance *d*_*j*_ away from the CpG undergoing the reaction. Thus, the sum occurs over distance functions *f*_*m*_(*d*) for all other CpGs in the system currently in the *m* state, scaled by the collaborative rate constant.

Similarly, the total rate for the h → u reaction becomes

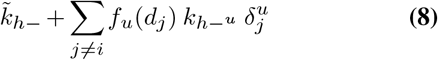

and so on.

### Coarse-Grained Model Approximation

We used a Chemical Master Equation (CME) framework to compute the steady-state distribution of methylation states, given a set of model parameters. We introduced a novel coarse-grained approximation of the full model that enables rapid computation of the methylation landscape, and is thus amenable to fitting parameters to data. Our approach is related to a recent mean-field method introduced by (78). Whereas their method is focused mainly on nearest-neighbor interactions, our method uses a distance function to compute interactions and incorporates the natural variation in inter-CpG spacing. However, our model cannot distinguish behavior of CpGs in a cluster, e.g. in the center versus the edge. Briefly, for a system of *N* CpGs at genomic positions {*s*_1_, *s*_2_, *s*_3_, …, *s*_*N*_}, rather than enumerate all possible 3^*N*^ system states, the system is described by its “macrostate”, {*N*_*u*_, *N*_*h*_, *N*_*m*_}, the number of CpGs in each of the three states, where *N*_*u*_ + *N*_*h*_ + *N*_*m*_ = *N*. Then the probability of an individual CpG being in *u, h, m*, given macrostate *r*, is {*p*_*u*|*r*_, *p*_*h*|*r*_, *p*_*m*|*r*_} = {*N*_*u*_, *N*_*h*_, *N*_*m r*_*/N*}. The average collaborative demethylating “field” in the *r*th macrostate is given by

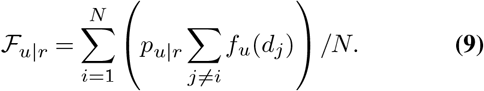

Similarly, the average collaborative methylating field in the *r*th macrostate is

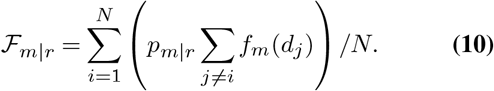

Finally, the chemical reaction network model is constructed as

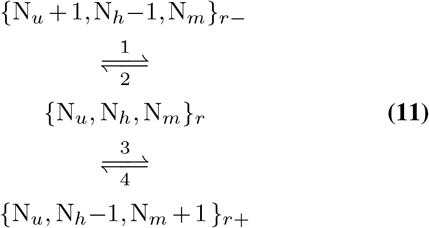

with the reaction rates given by

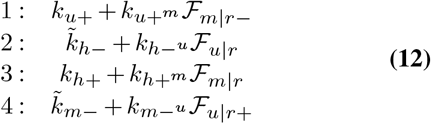

For a system of *N* CpGs, the number of macrostates is given by the triangular number

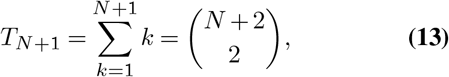

which increases much more slowly than 3^*N*^, making it possible to calculate distributions for larger numbers of CpGs. (Computations are tractable when the number of states is *<* ≈ 10^4^, limiting the system to ≤ 8 CpGs for the full model and up to ≈ 140 CpGs for the approximate, macrostate model).

### Fitting Model to Data

The chemical system can be expressed as 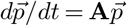, where 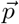 is the *T*_*N*+1_ ×1 vector holding probabilities for the group to be found in each possible macrostate, and ***A*** is the *T*_*N*+1_ × *T*_*N*+1_ reaction rate matrix, holding reaction rate constants connecting each macrostate as in Eq.1. The steady-state probabilities over each methylation macrostate for a given set of parameters were computed using MATLAB’s eigenvalue solver. To compute hypo/inter/hyper vs. CpG-density curves for each trial set of parameters, the simulated system density was varied by fixing *N* = 27 CpGs and varying the inter-CpG distance {*d, d* = 2, 3, 4, 5, …100}. The computed probability vector is mapped onto the Hyper/Inter/Hypo classification according to the methylation ratio of each macrostate, given by (0.5*N*_*h*_ + *N*_*m*_)*/N*. (That is, *m* are given a value of 1, *h* of 1/2, and *u* of 0.) The MATLAB optimization routine fmincon was used to optimize parameters. The representative parameter set fit to HUES8WT in Figs.6,7 is 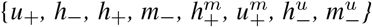 = {0.1, 0.1, 4, 0.298, 5.45, 0.155, 0.052, 18.8} (units h^−1^), {λ_*m*_, λ_*u*_*}* = {50, 43 bp}. The cell cycle time *N*_*t*_ is assumed to be 24 h.

### Software Packages and Code Availability

Data was analyzed via a combination of Python, Jupyter Notebook, and the *Pandas* software package (79); model fitting and figure generation were performed in *Matlab 2023b*. Code is made publicly available on Github via https://github.com/kbonsu/Ultrasens_DNAMethylation.

## Supporting Information

**Table S1.** Expanded Collection of Publicly Available WGBS Datasets Considered. Note that Illumina HiSeq 2000 is used for the HUES64 WT/KO and IMR90 datasets, HiSeq 6000 for the HUES8 WT, and HiSeq 4000 for the HUES8-KOs.

| Cell Lineage | DNMT/TET-KO Type | GEO Accession No. |
| --- | --- | --- |
| HUES64 | WT | GSM1112841 |
| HUES64 | DNMT-3A (Early, P24+7) | GSM1545002 |
| HUES64 | DNMT-3B (Early, P24+6) | GSM1545003 |
| HUES64 | DNMT-3A (Late, P24+22) | GSM1545005 |
| HUES64 | DNMT-3B (Late, P24+22) | GSM1545006 |
| HUES64 | DNMT-3A+3B (DKO) (Early, P24+5+2) | GSM154004 |
| HUES64 | DNMT-3A+3B (DKO) (Late, P24+5+17) | GSM1545007 |
| HUES8 | WT | GSM3618718 |
| HUES8 | DNMT-3A+3B (DKO, P6) | GSM4458672 |
| HUES8 | TET1-3 (TKO) | GSM3618720 |
| HUES8 | TET1-3 + DNMT-3B (QKO) | GSM3618719 |
| HUES8 | TET1-3 + DNMT-3A+3B (PKO, P6) | GSM3662266 |
| IMR90 | WT | GSM1204464 |

**Fig. S1.**
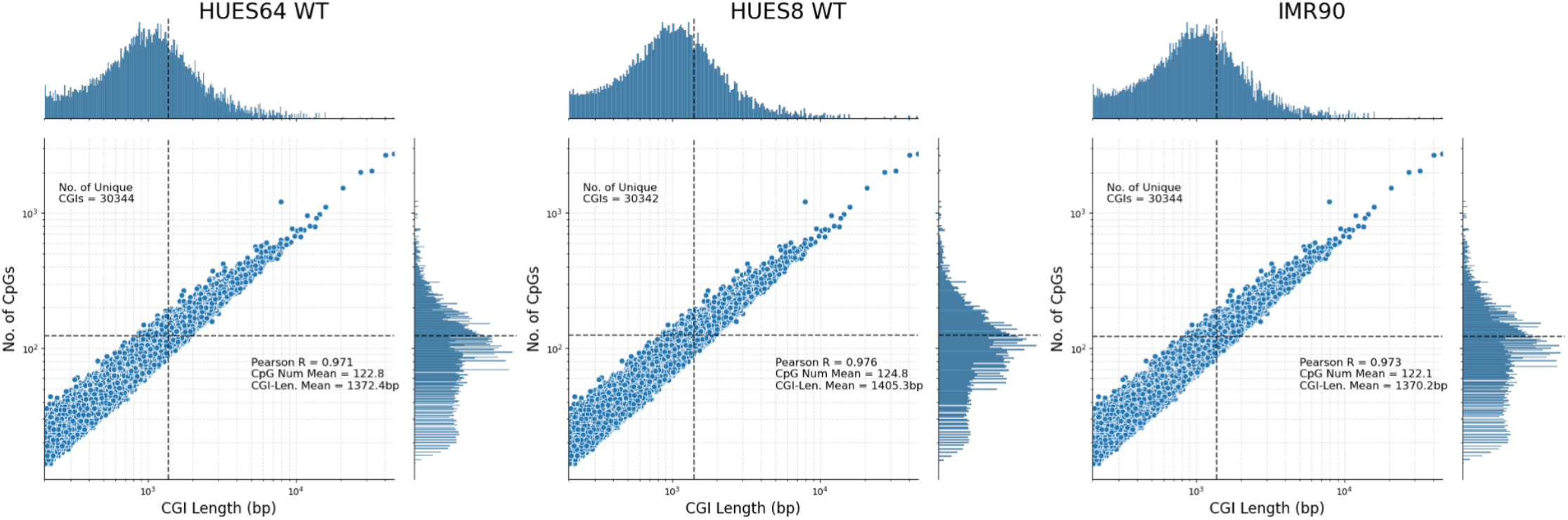
CpG Numbers of Annotated CpG Islands in Wild-Type HUES64, HUES8, and IMR90 Cell Lines are plotted as a function of CpG Island Length (bp), with external histograms displaying population counts of these metrics within the dataset, to illustrate linearly increasing relationship between CpG content and distance of these genomic features. Number of Unique Annotated CGIs, Pearson Correlation Coefficients, mean CpG Number and mean CGI length for each dataset.

**Fig. S2.**
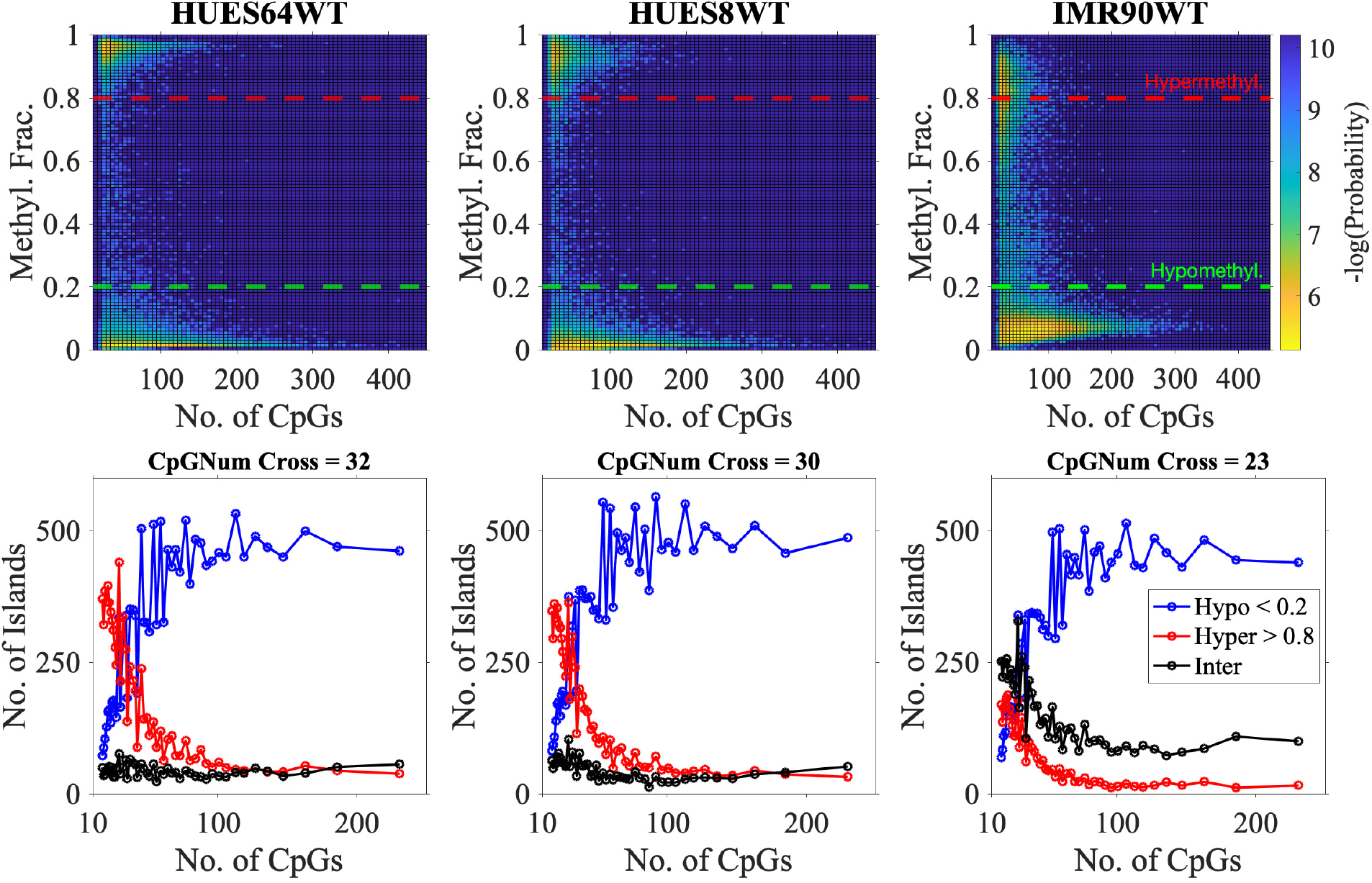
Dependence of CpG Island (CGI) methylation on CGI length in HUES64, HUES8, and IMR90 cells. *Top Row:* Bivariate heat maps of CGI-level averaging of bisulfite sequencing data comparing associated Methyl Fraction and Island Length; color corresponds to negative log-probability across binned tiles (yellow corresponds to highest density of islands). For each island, Methylation Fraction is defined as the sum over CpG-methylation values, divided by the number of CpGs in the island. CGI length is defined as the distance in bp between the first and last cytosine in CpGs associated to the island. *Bottom Row:* Curves display the number of CGIs that are classified into *Hypomethylated* (<20% Methylation Fraction, Dotted Blue Threshold), *Hypermethylated* (>80% Methylation Fraction, Dotted Red Threshold), and Intermediate methylation states, as a function of length. Length bins are determined by percentiles, giving an approximately equal number of islands in each length-bin. For each cell type, EL50 gives the length in bp at which the red and blue curves cross; that is, the length at which the number of islands classified into Hypo- and Hypermethylated groups is equal.

**Fig. S3.**
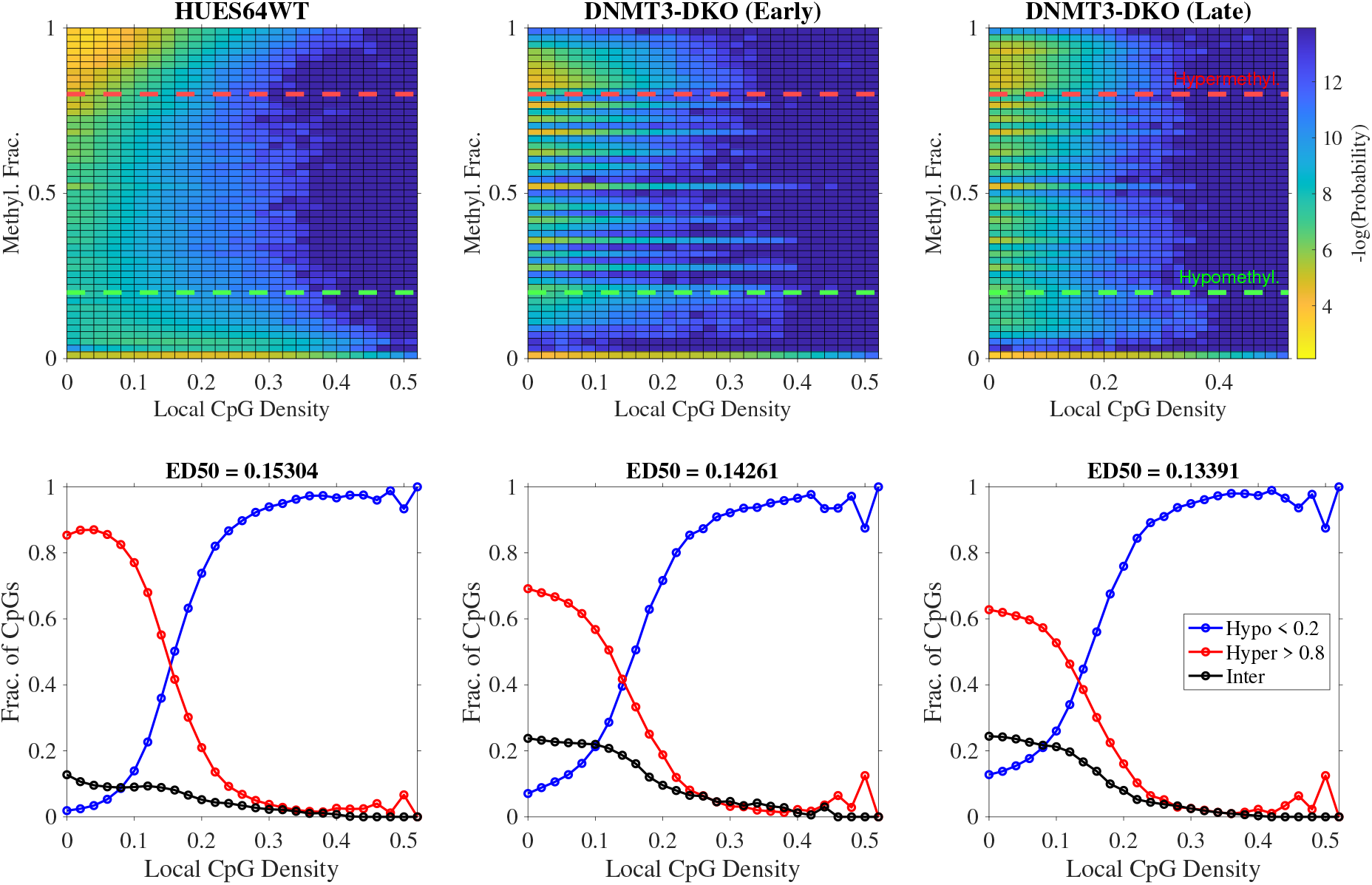
Bivariate Histograms of Individual CpG-level Methylation with respect to a window size of 50bp (up/downstream) recapitulate bimodal behavior seen at the broader, CGI-level in HUES64 DNMT3A/B DKO System. Top Rows show 2D Heat Maps Individual CpG-level of CpG bisulfite data comparing associated Methyl Fraction and Local CpG density; color corresponds to negative log-probability across binned tiles. Lower row display curves in which CpGs are sorted into Hypomethylated, Hypermethylated, and Intermediate methylation states as described in the main text.

**Fig. S4.**
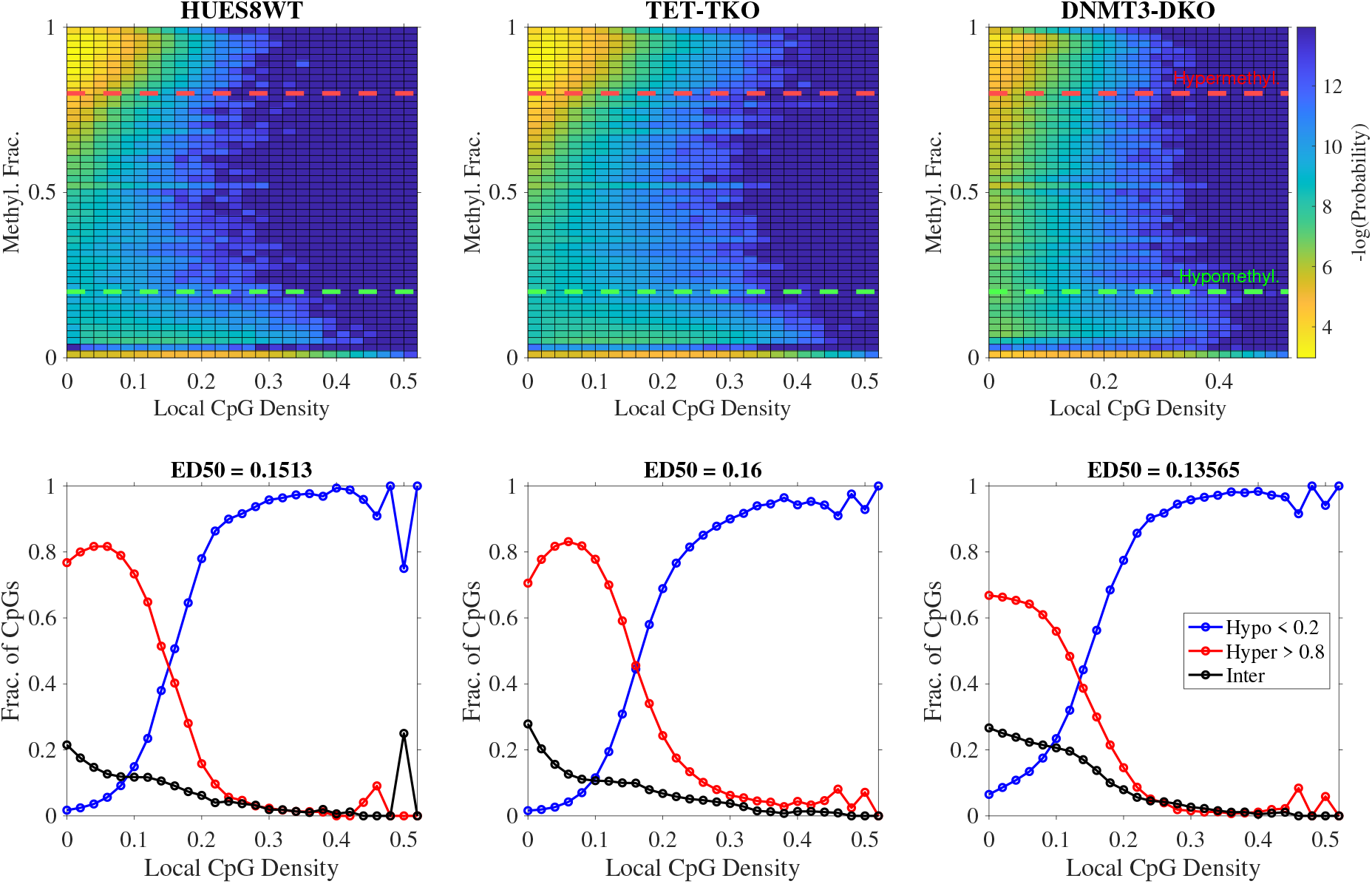
Bivariate Histograms of Individual CpG-level Methylation with respect to a window size of 50bp (up/downstream) recapitulate bimodal behavior seen at the broader, CGI-level in HUES8 TET1-3 TKO and DNMT3A/B DKO Systems. Top Rows show 2D Heat Maps Individual CpG-level of CpG bisulfite data comparing associated Methyl Fraction and Local CpG density; color corresponds to negative log-probability across binned tiles. Lower row display curves in which CpGs are sorted into Hypomethylated, Hypermethylated, and Intermediate methylation states as described in the main text.

**Fig. S5.**
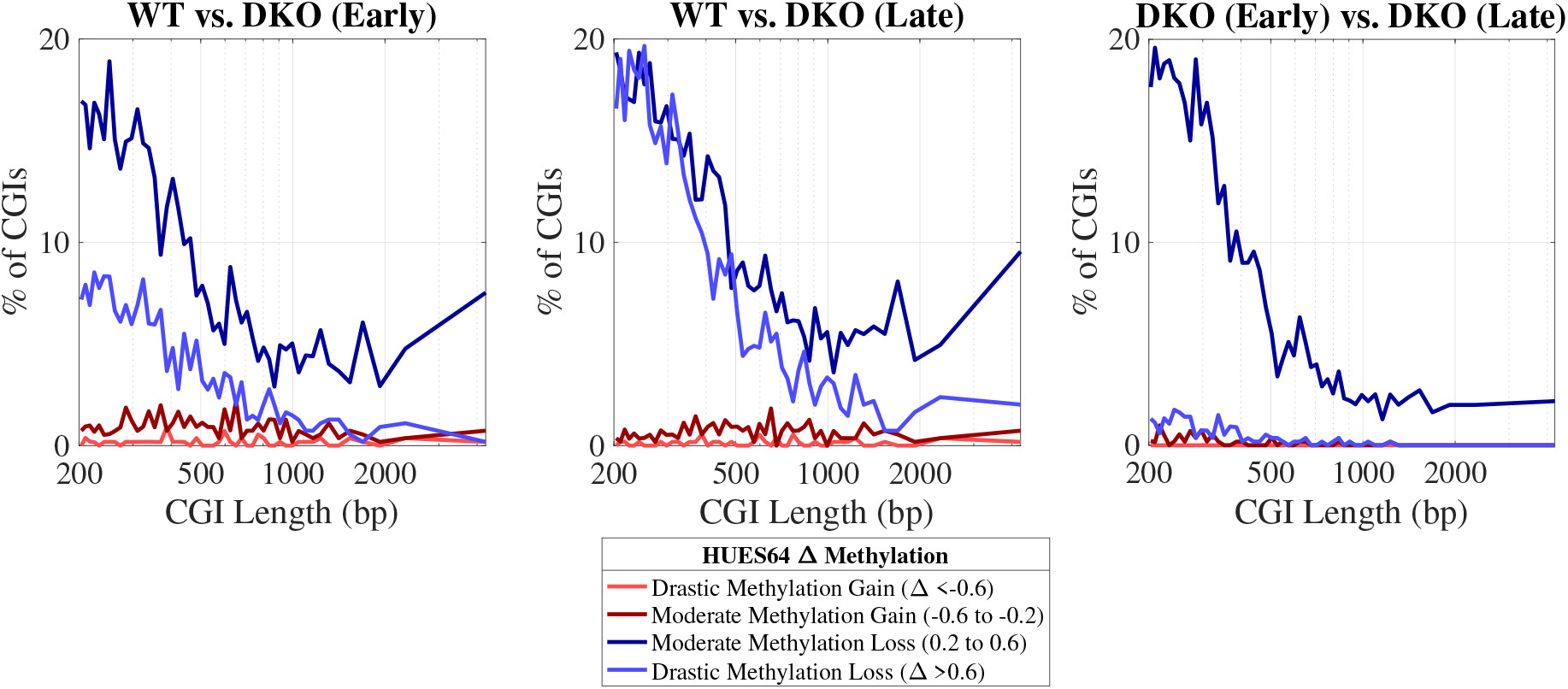
Quantification of State-Switching CGIs in HUES64 DNMT3-DKO system. Panels calculate percentage of CGIs which experienced marked changes given the annotated cell types; islands which experienced less than a 20% change in methylation are omitted. In all shown comparisons, drastic and moderate methylation changes are associated with shorter CGIs. When comparing Early vs. Late timepoints, longer CGIs experience less moderate methylation loss, suggesting these islands are less susceptible to time-dependent changes of methylation due to loss of de novo methyltransferase activity.

**Fig. S6.**
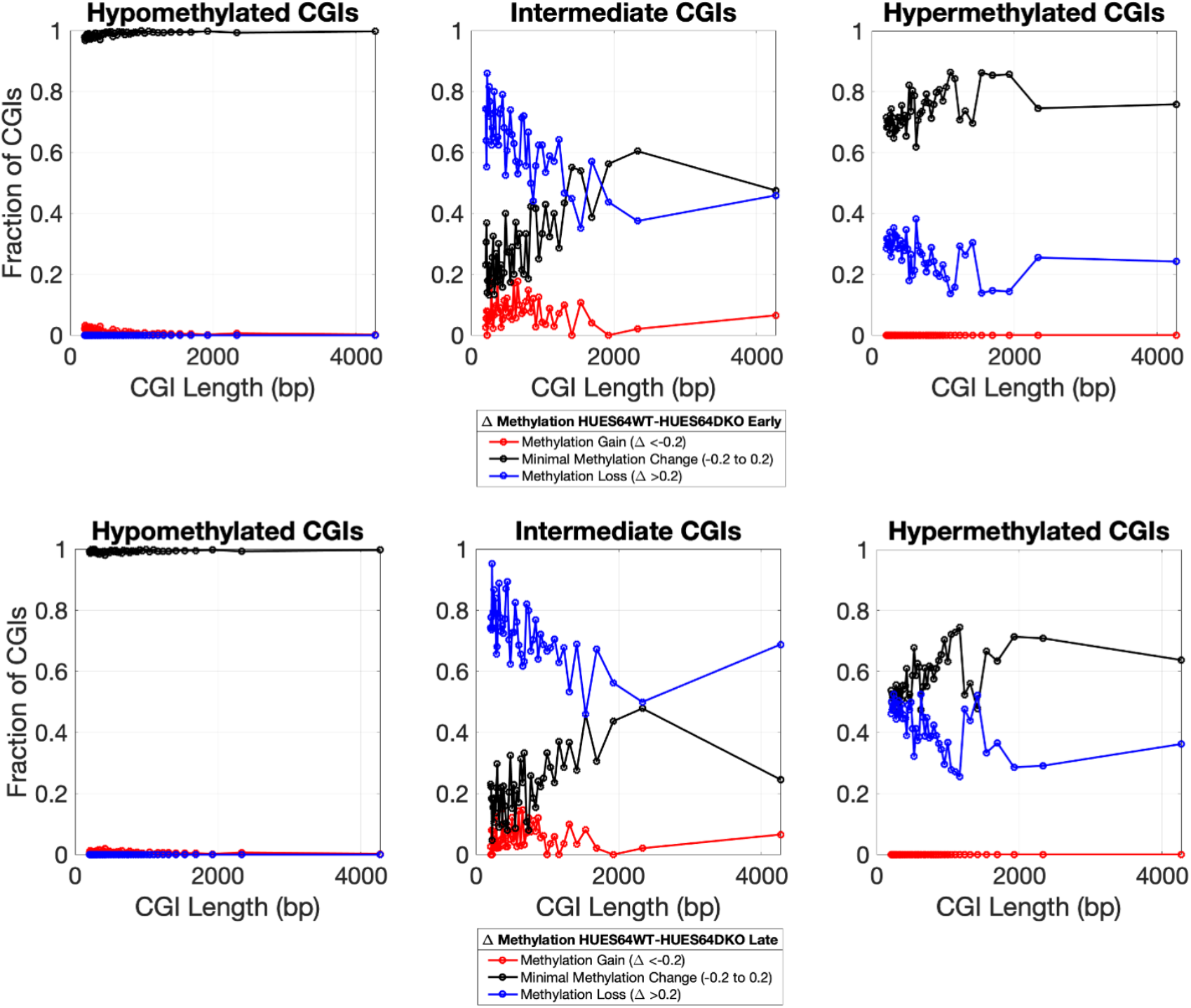
Pre-Classifying State-Switching CGIs in HUES64 DNMT3A/B DKO System. CGIs are pre-classified into their Hypomethyltated, Intermediate, or Hypermethylated macrostates, before calculating percentage of CGIs which experienced marked changes in DKO Early (*Top Row*) and Late (*Bottom Row*) timepoints compared to the WT; islands which experienced less than a 20% change in methylation are omitted.

**Fig. S7.**
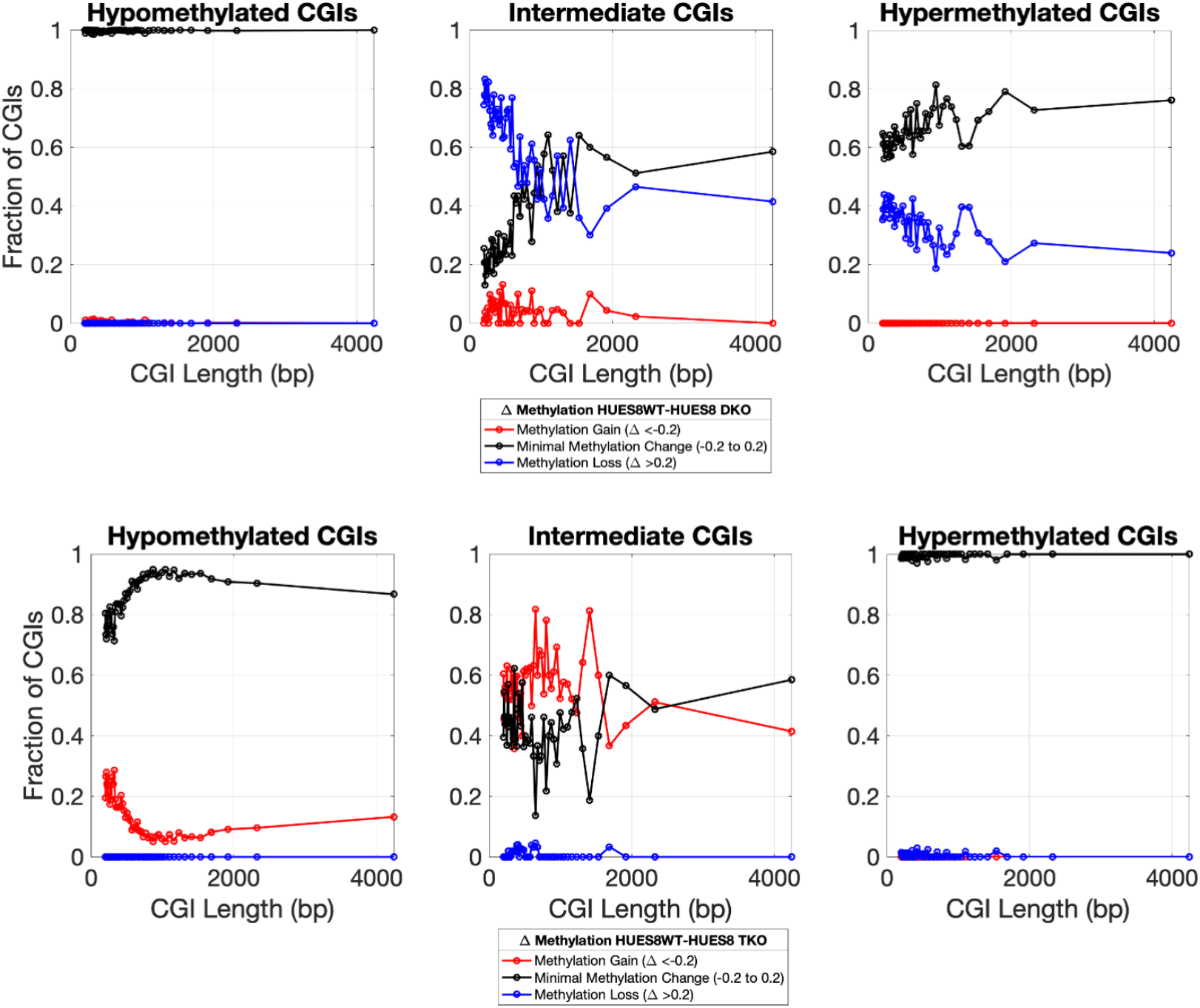
Pre-Classifying State-Switching CGIs in HUES8 TET1-3 TKO and DNMT3A/B DKO System. CGIs are pre-classified into their Hypomethyltated, Intermediate, or Hypermethylated macrostates, before calculating percentage of CGIs which experienced marked changes in DKO (*Top Row*) and TKO (*Bottom Row*) knockout systems compared to the WT; islands which experienced less than a 20% change in methylation are omitted.

**Fig. S8.**
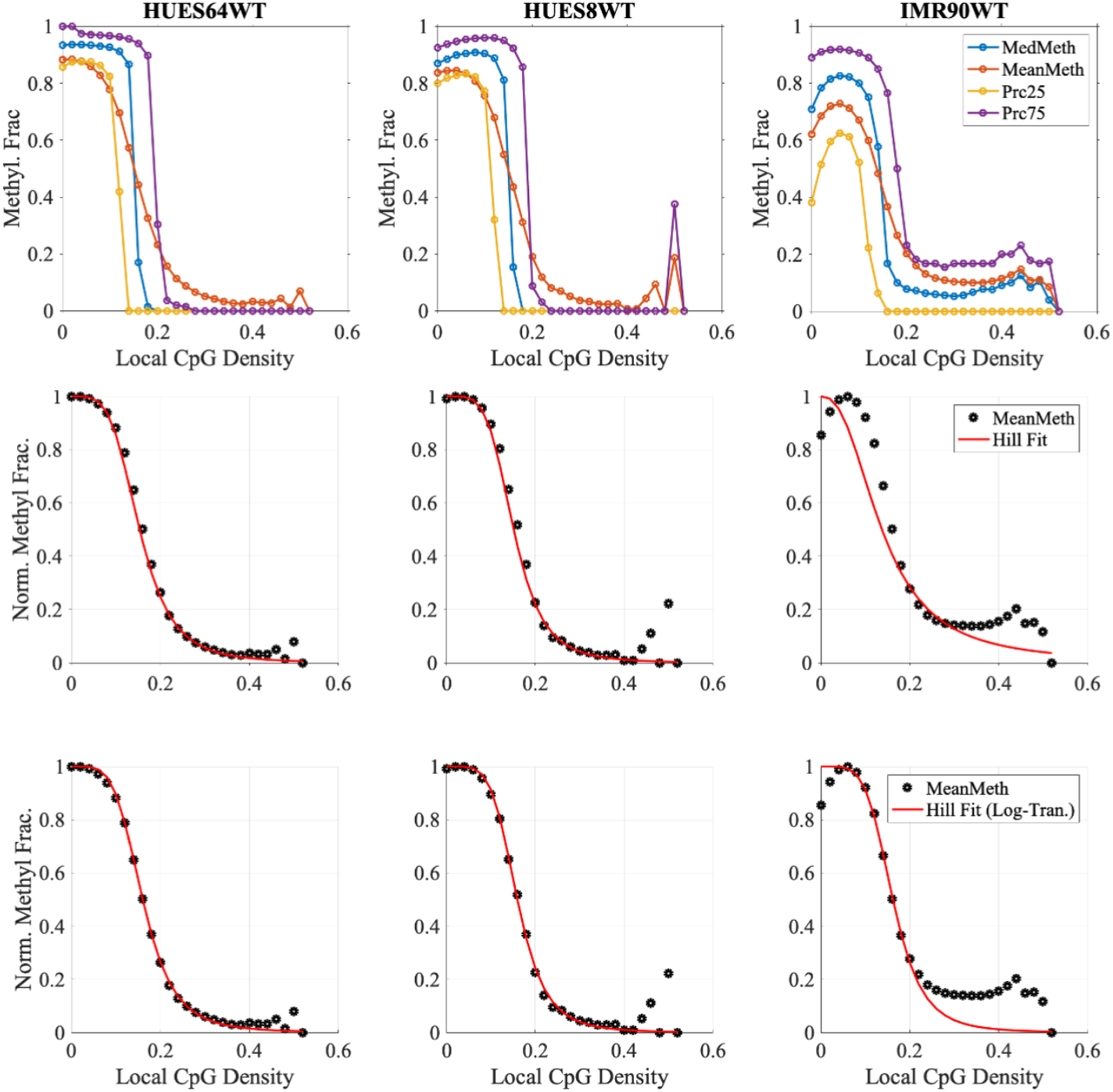
(*Row 1*): Median, Mean, and 25th-/75th-percentiles of Individual CpGs are plotted as a function of local CpG density, as described prior, in *wild-type* HUES64, HUES8, and IMR90 cell lines. Mean methylation curves are fitted to Hill Function equations directly, by minimizing sum of squared differences to the entire landscape (*Simple, Row 2*) and by fitting the steepest part of the curve after log-transforming the data (*Row 3*).

**Fig. S9.**
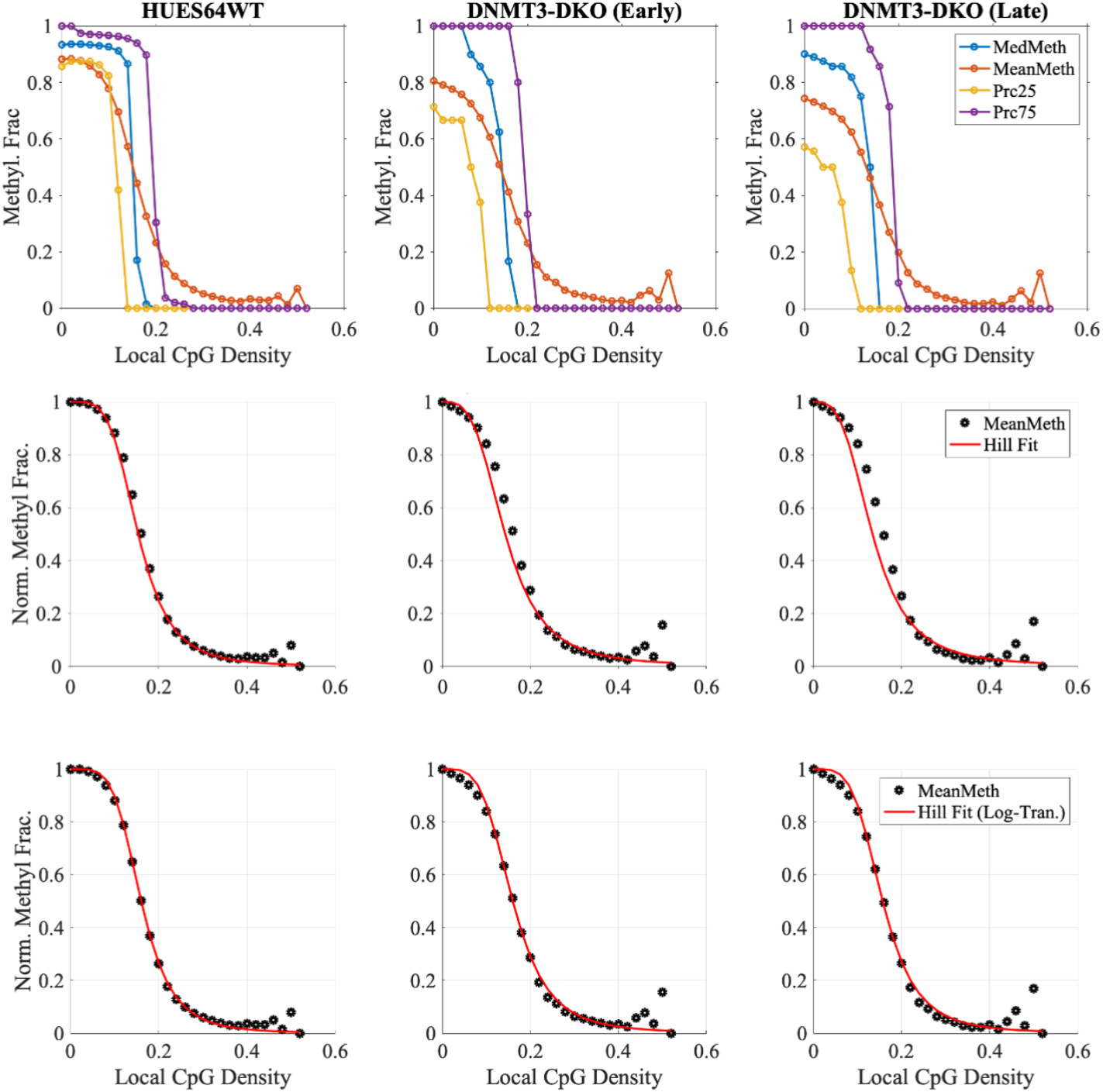
*Row 1*: Median, Mean, and 25th-/75th-percentiles of Individual CpGs are plotted as a function of local CpG density, as described prior, in *wild-type* HUES64 DKO systems. Mean methylation curves are fitted to Hill Function equations directly, by minimizing sum of squared differences to the entire landscape (*Simple, Row 2*) and by fitting the steepest part of the curve after log-transforming the data (*Row 3*).

**Fig. S10.**
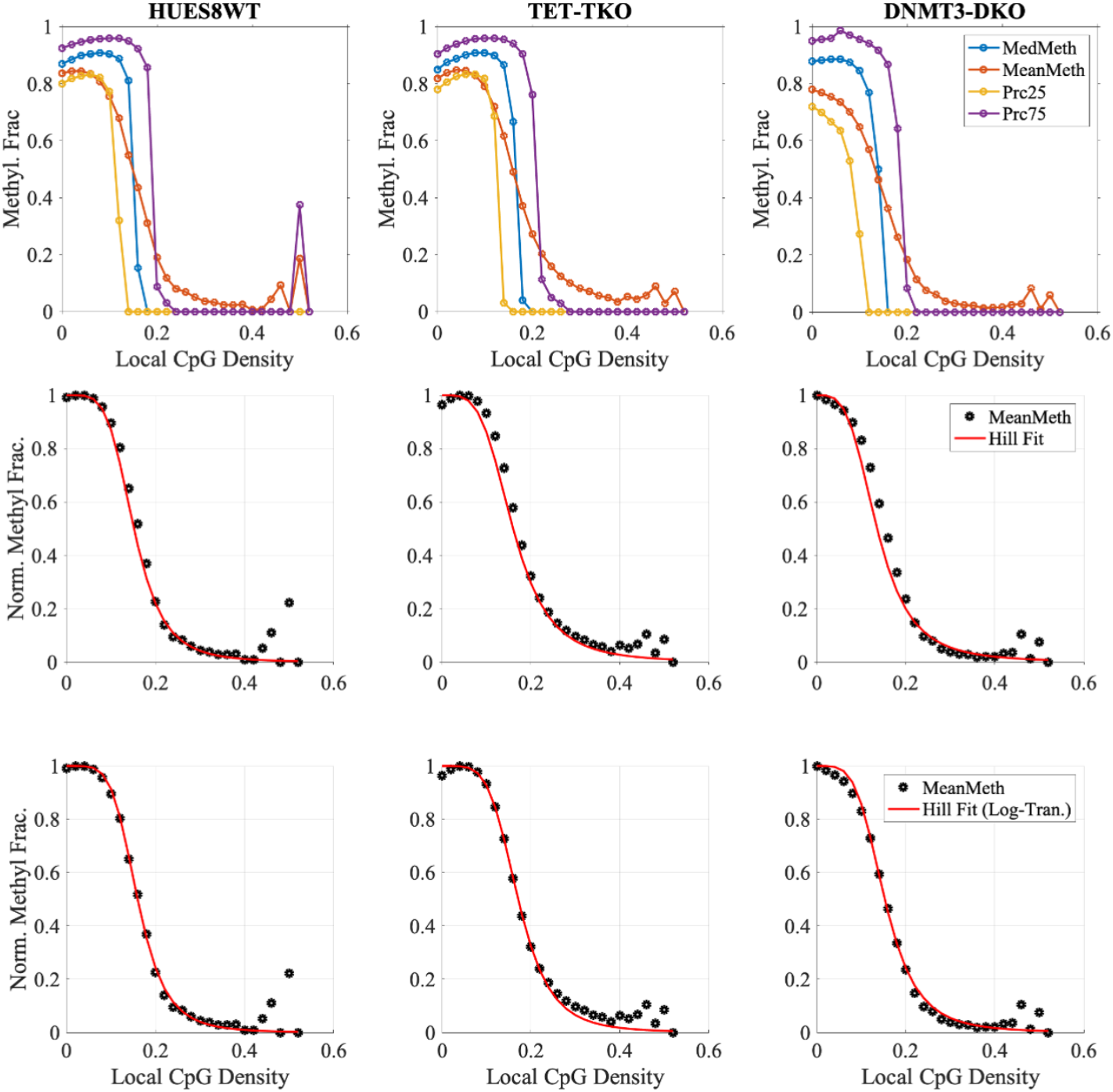
*Row 1*: Median, Mean, and 25th-/75th-percentiles of Individual CpGs are plotted as a function of local CpG density, as described prior, in *wild-type* HUES8 DKO and TKO systems. Mean methylation curves are fitted to Hill Function equations directly, by minimizing sum of squared differences to the entire landscape (*Simple, Row 2*) and by fitting the steepest part of the curve after log-transforming the data (*Row 3*).

**Fig. S11.**
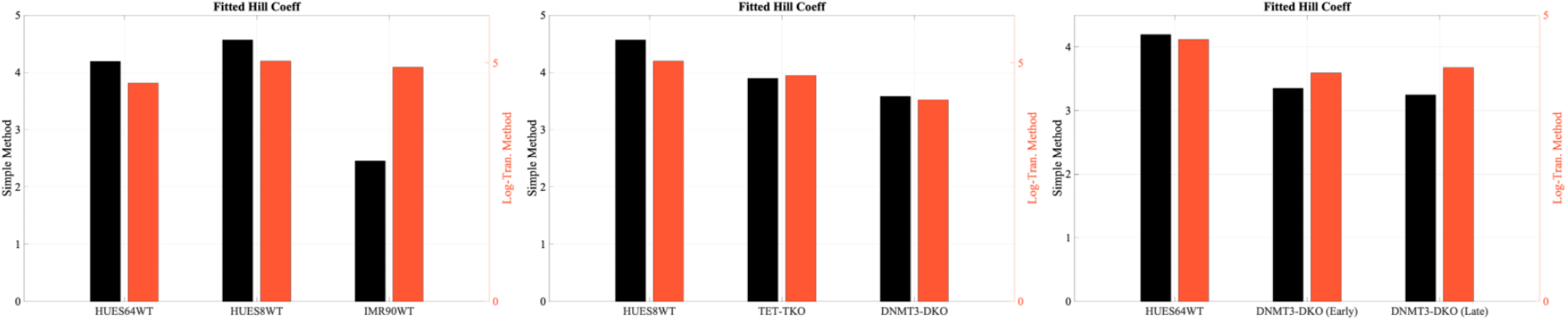
Hill function exponents derived from both the simple fitting method (*Left bars*) and via log-transforming the data to fit the steepest portion of the curve (*Right bars*); panels compare *wild-type* cells (*Left*), HUES64 DKO cells (*Center*), and HUES8 TKO/DKO cells (*Right*).

## Notes

### Competing Interest Statement

The authors have declared no competing interest.

## Bibliography

1. Shiv IS Grewal and Danesh Moazed. Heterochromatin and epigenetic control of gene expression. Science, 301(5634):798–802, August 2003.

2. Shikhar Sharma, Theresa K. Kelly, and Peter A. Jones. Epigenetics in cancer. Carcinogenesis, 31(1):27–36, January 2010. ISSN 0143-3334. doi: 10.1093/carcin/bgp220. Number:

3. Jacob L. Glass, Reid F. Thompson, Batbayar Khulan, Maria E. Figueroa, Emmanuel N. Olivier, Erin J. Oakley, Gary Van Zant, Eric E. Bouhassira, Ari Melnick, Aaron Golden, Melissa J. Fazzari, and John M. Greally. CG dinucleotide clustering is a species-specific property of the genome. Nucleic Acids Research, 35(20):6798–6807, October 2007. doi: 10.1093/nar/gkm489.

4. M. Gardiner-Garden and M. Frommer. CpG islands in vertebrate genomes. Journal of Molecular Biology, 196(2):261–282, July 1987. doi: 10.1016/0022-2836(87)90689-9.

5. Leng Han, Bing Su, Wen-Hsiung Li, and Zhongming Zhao. CpG island density and its correlations with genomic features in mammalian genomes. Genome Biology, 9(5):R79, 2008. doi: 10.1186/gb-2008-9-5-r79.

6. Gulsah Altun, Jeanne F. Loring, and Louise C. Laurent. DNA methylation in embryonic stem cells. Journal of Cellular Biochemistry, 109(1):1–6, January 2010. ISSN 0730-2312. doi: 10.1002/jcb.22374. Number: 1.

7. Serge Saxonov, Paul Berg, and Douglas L Brutlag. A genome-wide analysis of CpG dinucleotides in the human genome distinguishes two distinct classes of promoters. Proc. Natl. Acad. Sci. U. S. A., 103(5):1412–1417, January 2006.

8. Maria Gutierrez-Arcelus, Tuuli Lappalainen, Stephen B Montgomery, Alfonso Buil, Halit Ongen, Alisa Yurovsky, Julien Bryois, Thomas Giger, Luciana Romano, Alexandra Planchon, Emilie Falconnet, Deborah Bielser, Maryline Gagnebin, Ismael Padioleau, Christelle Borel, Audrey Letourneau, Periklis Makrythanasis, Michel Guipponi, Corinne Gehrig, Stylianos E Antonarakis, and Emmanouil T Dermitzakis. Passive and active DNA methylation and the interplay with genetic variation in gene regulation. Elife, 2:e00523, June 2013.

9. Wanding Zhou, Gangning Liang, Peter L Molloy, and Peter A Jones. DNA methylation enables transposable element-driven genome expansion. Proc. Natl. Acad. Sci. U. S. A., 117(32):19359–19366, August 2020.

10. Atsuya Nishiyama, Luna Yamaguchi, Jafar Sharif, Yoshikazu Johmura, Takeshi Kawamura, Keiko Nakanishi, Shintaro Shimamura, Kyohei Arita, Tatsuhiko Kodama, Fuyuki Ishikawa, Haruhiko Koseki, and Makoto Nakanishi. Uhrf1-dependent H3K23 ubiquitylation couples maintenance DNA methylation and replication. Nature, 502(7470):249–253, October 2013. ISSN 1476-4687. doi: 10.1038/nature12488. Number: 7470 Publisher: Nature Publishing Group.

11. Harsimran Sidhu and Neena Capalash. UHRF1: The key regulator of epigenetics and molecular target for cancer therapeutics. Tumour Biol., 39(2):1010428317692205, February 2017.

12. Christian Bronner, Mahmoud Alhosin, Ali Hamiche, and Marc Mousli. Coordinated dialogue between UHRF1 and DNMT1 to ensure faithful inheritance of methylated DNA patterns. Genes, 10(1):65, January 2019. doi: 10.3390/genes10010065.

13. Jing Liao, Rahul Karnik, Hongcang Gu, Michael J. Ziller, Kendell Clement, Alexander M. Tsankov, Veronika Akopian, Casey A. Gifford, Julie Donaghey, Christina Galonska, Ramona Pop, Deepak Reyon, Shengdar Q. Tsai, William Mallard, J. Keith Joung, John L. Rinn, Andreas Gnirke, and Alexander Meissner. Targeted disruption of DNMT1, DNMT3A and DNMT3B in human embryonic stem cells. Nature genetics, 47(5):469–478, May 2015. ISSN 1061-4036. doi: 10.1038/ng.3258. Number: 5.

14. Frederic Chedin, Michael R Lieber, and Chih-Lin Hsieh. The DNA methyltransferase-like protein DNMT3L stimulates de novo methylation by dnmt3a. Proc. Natl. Acad. Sci. U. S. A., 99(26):16916–16921, December 2002.

15. Christopher E Duymich, Jessica Charlet, Xiaojing Yang, Peter A Jones, and Gangning Liang. DNMT3B isoforms without catalytic activity stimulate gene body methylation as accessory proteins in somatic cells. Nat. Commun., 7:11453, April 2016.

16. Nathan R Rose and Robert J Klose. Understanding the relationship between DNA methylation and histone lysine methylation. Biochim. Biophys. Acta, 1839(12):1362–1372, December 2014.

17. Kasper Dindler Rasmussen and Kristian Helin. Role of TET enzymes in DNA methylation, development, and cancer. Genes & Development, 30(7):733–750, April 2016. ISSN 0890-9369. doi: 10.1101/gad.276568.115. Number: 7.

18. Tao Zhu, Anthony P Brown, and Hong Ji. The emerging role of ten-eleven translocation 1 in epigenetic responses to environmental exposures. Epigenetics Insights, 13: 2516865720910155, March 2020.

19. Ming Zhang, Jian Wang, Kaixiang Zhang, Guozhen Lu, Yuming Liu, Keke Ren, Wenting Wang, Dazhuan Xin, Lingli Xu, Honghui Mao, Junlin Xing, Xingchun Gao, Weilin Jin, Kalen Berry, Katsuhiko Mikoshiba, Shengxi Wu, Q Richard Lu, and Xianghui Zhao. Ten-eleven translocation 1 mediated-DNA hydroxymethylation is required for myelination and remyelination in the mouse brain. Nat. Commun., 12(1):5091, August 2021.

20. Jocelyn Charlton, Eunmi J Jung, Alexandra L Mattei, Nina Bailly, Jing Liao, Eric J Martin, Pay Giesselmann, Björn Brändl, Elena K Stamenova, Franz-Josef Müller, Evangelos Kiskinis, Andreas Gnirke, Zachary D Smith, and Alexander Meissner. TETs compete with DNMT3 activity in pluripotent cells at thousands of methylated somatic enhancers. Nat. Genet., 52(8):819–827, August 2020.

21. Fushun Chen, Qingzheng Zhang, Xiaodi Deng, Xia Zhang, Chengjun Chen, Dekang Lv, Yulong Li, Dan Li, Yu Zhang, Peiying Li, et al. Conflicts of cpg density and dna methylation are proximally and distally involved in gene regulation in human and mouse tissues. Epigenetics, 13(7):721–741, 2018.

22. Dimos Gaidatzis, Lukas Burger, Rabih Murr, Anita Lerch, Sophie Dessus-Babus, Dirk Schübeler, and Michael B Stadler. Dna sequence explains seemingly disordered methylation levels in partially methylated domains of mammalian genomes. PLoS genetics, 10(2):e1004143, 2014.

23. Wanding Zhou, Huy Q Dinh, Zachary Ramjan, Daniel J Weisenberger, Charles M Nicolet, Hui Shen, Peter W Laird, and Benjamin P Berman. Dna methylation loss in late-replicating domains is linked to mitotic cell division. Nature genetics, 50(4):591–602, 2018.

24. John R Edwards, Anne H O’Donnell, Robert A Rollins, Heather E Peckham, Clarence Lee, Maria H Milekic, Benjamin Chanrion, Yutao Fu, Tao Su, Hanina Hibshoosh, et al. Chromatin and sequence features that define the fine and gross structure of genomic methylation patterns. Genome research, 20(7):972–980, 2010.

25. Cecilia Lövkvist, Ian B. Dodd, Kim Sneppen, and Jan O. Haerter. DNA methylation in human epigenomes depends on local topology of CpG sites. Nucleic Acids Research, 44(11): 5123–5132, June 2016. ISSN 0305-1048, 1362-4962. doi: 10.1093/nar/gkw124.

26. Jan O. Haerter, Cecilia Lövkvist, Ian B. Dodd, and Kim Sneppen. Collaboration between CpG sites is needed for stable somatic inheritance of DNA methylation states. Nucleic Acids Research, 42(4):2235–2244, February 2014. ISSN 1362-4962, 0305-1048. doi: 10.1093/nar/gkt1235.

27. Jocelyn Charlton, Eunmi J Jung, Alexandra L Mattei, Nina Bailly, Jing Liao, Eric J Martin, Pay Giesselmann, Björn Brändl, Elena K Stamenova, Franz-Josef Müller, et al. Tets compete with dnmt3 activity in pluripotent cells at thousands of methylated somatic enhancers. Nature genetics, 52(8):819–827, 2020.

28. Albert Goldbeter and Daniel E Koshland Jr. An amplified sensitivity arising from covalent modification in biological systems. Proceedings of the National Academy of Sciences, 78 (11):6840–6844, 1981.

29. Howard Cedar. Dna methylation and gene activity. Cell, 53(1):3–4, April 1988. ISSN 0092-8674. doi: 10.1016/0092-8674(88)90479-5.

30. James E. Ferrell and Sang Hoon Ha. Ultrasensitivity part i: Michaelian responses and zero-order ultrasensitivity. Trends in Biochemical Sciences, 39(10):496–503, October 2014. ISSN 0968-0004. doi: 10.1016/j.tibs.2014.08.003.

31. James E. Ferrell, Jr, and Sang Hoon Ha. Ultrasensitivity part ii: multisite phosphorylation, stoichiometric inhibitors, and positive feedback. Trends in Biochemical Sciences, 39(11): 556–569, November 2014. ISSN 0968-0004. doi: 10.1016/j.tibs.2014.09.003.

32. James E. Ferrell and Sang Hoon Ha. Ultrasensitivity part iii: cascades, bistable switches, and oscillators. Trends in Biochemical Sciences, 39(12):612–618, December 2014. ISSN 0968-0004. doi: 10.1016/j.tibs.2014.10.002.

33. Brian Munsky and Mustafa Khammash. The finite state projection algorithm for the solution of the chemical master equation. The Journal of chemical physics, 124(4), 2006.

34. Brian K Chu, Margaret J Tse, Royce R Sato, and Elizabeth L Read. Markov state models of gene regulatory networks. BMC systems biology, 11:1–17, 2017.

35. Timothy S. Gardner, Charles R. Cantor, and James J. Collins. Construction of a genetic toggle switch in escherichia coli. Nature, 403(6767):339–342, January 2000. ISSN 1476-4687. doi: 10.1038/35002131.

36. Anand Kumar Sharma, Radhika Khandelwal, and Christian Wolfrum. Futile cycles: Emerging utility from apparent futility. Cell Metabolism, April 2024. ISSN 1550-4131. doi: 10.1016/j.cmet.2024.03.008.

37. A. Cornish-Bowden. Fundamentals of Enzyme Kinetics. Wiley, 2013. ISBN 9783527665495.

38. David C LaPorte and Daniel E Koshland Jr. Phosphorylation of isocitrate dehydrogenase as a demonstration of enhanced sensitivity in covalent regulation. Nature, 305(5932):286–290, 1983.

39. Marilyn H Meinke, Jonathan S Bishop, and Ronald D Edstrom. Zero-order ultrasensitivity in the regulation of glycogen phosphorylase. Proceedings of the National Academy of Sciences, 83(9):2865–2868, 1986.

40. Gustavo J Melen, Sagi Levy, Naama Barkai, and Ben-Zion Shilo. Threshold responses to morphogen gradients by zero-order ultrasensitivity. Molecular systems biology, 1(1):2005– 0028, 2005.

41. Mohan K Malleshaiah, Vahid Shahrezaei, Peter S Swain, and Stephen W Michnick. The scaffold protein ste5 directly controls a switch-like mating decision in yeast. Nature, 465 (7294):101–105, 2010.

42. James E Ferrell and Sang Hoon Ha. Ultrasensitivity part i: Michaelian responses and zero-order ultrasensitivity. Trends in biochemical sciences, 39(10):496–503, 2014.

43. Michael Weber, Ines Hellmann, Michael B Stadler, Liliana Ramos, Svante Pääbo, Michael Rebhan, and Dirk Schübeler. Distribution, silencing potential and evolutionary impact of promoter dna methylation in the human genome. Nature genetics, 39(4):457–466, 2007.

44. Alexander Meissner, Tarjei S Mikkelsen, Hongcang Gu, Marius Wernig, Jacob Hanna, Andrey Sivachenko, Xiaolan Zhang, Bradley E Bernstein, Chad Nusbaum, David B Jaffe, et al. Genome-scale dna methylation maps of pluripotent and differentiated cells. Nature, 454 (7205):766–770, 2008.

45. Christoph Bock, Martina Paulsen, Sascha Tierling, Thomas Mikeska, Thomas Lengauer, and Jörn Walter. Cpg island methylation in human lymphocytes is highly correlated with dna sequence, repeats, and predicted dna structure. PLoS genetics, 2(3):e26, 2006.

46. Elisabeth Wachter, Timo Quante, Cara Merusi, Aleksandra Arczewska, Francis Stewart, Shaun Webb, and Adrian Bird. Synthetic cpg islands reveal dna sequence determinants of chromatin structure. Elife, 3:e03397, 2014.

47. Arnaud R Krebs, Sophie Dessus-Babus, Lukas Burger, and Dirk Schübeler. High-throughput engineering of a mammalian genome reveals building principles of methylation states at cg rich regions. Elife, 3:e04094, 2014.

48. Yuichi Mishima, Laura Brueckner, Saori Takahashi, Toru Kawakami, Junji Otani, Akira Shinohara, Kohei Takeshita, Ronald Garingalao Garvilles, Mikio Watanabe, Norio Sakai, et al. Enhanced processivity of dnmt1 by monoubiquitinated histone h3. Genes to cells, 25(1): 22–32, 2020.

49. Xuan Ming, Zhuqiang Zhang, Zhuoning Zou, Cong Lv, Qiang Dong, Qixiang He, Yangyang Yi, Yingfeng Li, Hailin Wang, and Bing Zhu. Kinetics and mechanisms of mitotic inheritance of dna methylation and their roles in aging-associated methylome deterioration. Cell research, 30(11):980–996, 2020.

50. Timothy H Bestor and Vernon M Ingram. Two dna methyltransferases from murine erythroleukemia cells: purification, sequence specificity, and mode of interaction with dna. Proceedings of the National Academy of Sciences, 80(18):5559–5563, 1983.

51. Giedrius Vilkaitis, Isao Suetake, Saulius Klimašauskas, and Shoji Tajima. Processive methylation of hemimethylated cpg sites by mouse dnmt1 dna methyltransferase. Journal of Biological Chemistry, 280(1):64–72, 2005.

52. Željko M Svedružić and Norbert O Reich. Mechanism of allosteric regulation of dnmt1’s processivity. Biochemistry, 44(45):14977–14988, 2005.

53. Andrea Hermann, Rachna Goyal, and Albert Jeltsch. The dnmt1 dna-(cytosine-c5)-methyltransferase methylates dna processively with high preference for hemimethylated target sites. Journal of Biological Chemistry, 279(46):48350–48359, 2004.

54. Rachna Goyal, Richard Reinhardt, and Albert Jeltsch. Accuracy of dna methylation pattern preservation by the dnmt1 methyltransferase. Nucleic acids research, 34(4):1182–1188, 2006.

55. Sabrina Adam, Hiwot Anteneh, Maximilian Hornisch, Vincent Wagner, Jiuwei Lu, Nicole E Radde, Pavel Bashtrykov, Jikui Song, and Albert Jeltsch. Dna sequence-dependent activity and base flipping mechanisms of dnmt1 regulate genome-wide dna methylation. Nature communications, 11(1):1–15, 2020.

56. Luis Busto-Moner, Julien Morival, Honglei Ren, Arjang Fahim, Zachary Reitz, Timothy L Downing, and Elizabeth L Read. Stochastic modeling reveals kinetic heterogeneity in postreplication DNA methylation. PLoS Comput. Biol., 16(4):e1007195, April 2020.

57. Honglei Ren, Robert B Taylor, Timothy L Downing, and Elizabeth L Read. Locally correlated kinetics of post-replication dna methylation reveals processivity and region specificity in dna methylation maintenance. Journal of the Royal Society Interface, 19(195):20220415, 2022.

58. Magnolia Bostick, Jong Kyong Kim, Pierre-Olivier Estève, Amander Clark, Sriharsa Pradhan, and Steven E Jacobsen. Uhrf1 plays a role in maintaining dna methylation in mammalian cells. Science, 317(5845):1760–1764, 2007.

59. Jafar Sharif, Masahiro Muto, Shin-ichiro Takebayashi, Isao Suetake, Akihiro Iwamatsu, Takaho A Endo, Jun Shinga, Yoko Mizutani-Koseki, Tetsuro Toyoda, Kunihiro Okamura, et al. The sra protein np95 mediates epigenetic inheritance by recruiting dnmt1 to methylated dna. Nature, 450(7171):908–912, 2007.

60. Xiaoli Liu, Qinqin Gao, Pishun Li, Qian Zhao, Jiqin Zhang, Jiwen Li, Haruhiko Koseki, and Jiemin Wong. Uhrf1 targets dnmt1 for dna methylation through cooperative binding of hemi-methylated dna and methylated h3k9. Nature communications, 4(1):1–13, 2013.

61. Xiwen Xing, Shinsuke Sato, Nai-Kei Wong, Kumi Hidaka, Hiroshi Sugiyama, and Masayuki Endo. Direct observation and analysis of tet-mediated oxidation processes in a dna origami nanochip. Nucleic Acids Research, 48(8):4041–4051, 2020.

62. Esta Tamanaha, Shengxi Guan, Katherine Marks, and Lana Saleh. Distributive processing by the iron (ii)/–-ketoglutarate-dependent catalytic domains of the tet enzymes is consistent with epigenetic roles for oxidized 5-methylcytosine bases. Journal of the American Chemical Society, 138(30):9345–9348, 2016.

63. Yufei Xu, Feizhen Wu, Li Tan, Lingchun Kong, Lijun Xiong, Jie Deng, Andrew J Barbera, Lijuan Zheng, Haikuo Zhang, Stephen Huang, et al. Genome-wide regulation of 5hmc, 5mc, and gene expression by tet1 hydroxylase in mouse embryonic stem cells. Molecular cell, 42(4):451–464, 2011.

64. Yufei Xu, Chao Xu, Akiko Kato, Wolfram Tempel, Jose Garcia Abreu, Chuanbing Bian, Yeguang Hu, Di Hu, Bin Zhao, Tanja Cerovina, et al. Tet3 cxxc domain and dioxygenase activity cooperatively regulate key genes for xenopus eye and neural development. Cell, 151(6):1200–1213, 2012.

65. Hao Wu, Ana C D’Alessio, Shinsuke Ito, Kai Xia, Zhibin Wang, Kairong Cui, Keji Zhao, Yi Eve Sun, and Yi Zhang. Dual functions of tet1 in transcriptional regulation in mouse embryonic stem cells. Nature, 473(7347):389–393, 2011.

66. Kristine Williams, Jesper Christensen, Marianne Terndrup Pedersen, Jens V Johansen, Paul AC Cloos, Juri Rappsilber, and Kristian Helin. Tet1 and hydroxymethylcytosine in transcription and dna methylation fidelity. Nature, 473(7347):343–348, 2011.

67. Charalampos Kyriakopoulos, Karl Nordström, Paula Linh Kramer, Judith Yumiko Gottfreund, Abdulrahman Salhab, Julia Arand, Fabian Müller, Ferdinand von Meyenn, Gabriella Ficz, Wolf Reik, et al. A comprehensive approach for genome-wide efficiency profiling of dna modifying enzymes. Cell reports methods, 2(3), 2022.

68. Paul Adrian Ginno, Dimos Gaidatzis, Angelika Feldmann, Leslie Hoerner, Dilek Imanci, Lukas Burger, Frederic Zilbermann, Antoine HFM Peters, Frank Edenhofer, Sébastien A Smallwood, et al. A genome-scale map of dna methylation turnover identifies site-specific dependencies of dnmt and tet activity. Nature communications, 11(1):2680, 2020.

69. Luis R Nassar, Galt P Barber, Anna Benet-Pagès, Jonathan Casper, Hiram Clawson, Mark Diekhans, Clay Fischer, Jairo Navarro Gonzalez, Angie S Hinrichs, Brian T Lee, Christopher M Lee, Pranav Muthuraman, Beagan Nguy, Tiana Pereira, Parisa Nejad, Gerardo Perez, Brian J Raney, Daniel Schmelter, Matthew L Speir, Brittney D Wick, Ann S Zweig, David Haussler, Robert M Kuhn, Maximilian Haeussler, and W James Kent. The ucsc genome browser database: 2023 update. Nucleic Acids Research, 51(D1):D1188–D1195, November 2022. ISSN 1362-4962. doi: 10.1093/nar/gkac1072.

70. Mehrnaz Fatemi, Andrea Hermann, Shriharsa Pradhan, and Albert Jeltsch. The activity of the murine dna methyltransferase dnmt1 is controlled by interaction of the catalytic domain with the n-terminal part of the enzyme leading to an allosteric activation of the enzyme after binding to methylated dna. Journal of molecular biology, 309(5):1189–1199, 2001.

71. Albert Jeltsch and Renata Z Jurkowska. New concepts in dna methylation. Trends in biochemical sciences, 39(7):310–318, 2014.

72. Rahul M Kohli and Yi Zhang. Tet enzymes, tdg and the dynamics of dna demethylation. Nature, 502(7472):472–479, 2013.

73. Sarah P Otto and Virginia Walbot. Dna methylation in eukaryotes: kinetics of demethylation and de novo methylation during the life cycle. Genetics, 124(2):429–437, 1990.

74. Arthur D Riggs. X inactivation, differentiation, and dna methylation. Cytogenetic and Genome Research, 14(1):9–25, 1975.

75. Laura B Sontag, Matthew C Lorincz, and E Georg Luebeck. Dynamics, stability and inheritance of somatic dna methylation imprints. Journal of theoretical biology, 242(4):890–899, 2006.

76. You Song, Honglei Ren, and Jinzhi Lei. Collaborations between cpg sites in dna methylation. International Journal of Modern Physics B, 31(20):1750243, 2017.

77. Loukas Zagkos, Mark Mc Auley, Jason Roberts, and Nikos I Kavallaris. Mathematical models of dna methylation dynamics: Implications for health and ageing. Journal of theoretical biology, 462:184–193, 2019.

78. Lyndsay Kerr, Duncan Sproul, and Ramon Grima. Cluster mean-field theory accurately predicts statistical properties of large-scale dna methylation patterns. Journal of the Royal Society Interface, 19(186):20210707, 2022.

79. Wes McKinney et al. Data structures for statistical computing in python. In Proceedings of the 9th Python in Science Conference, volume 445, pages 51–56. Austin, TX, 2010.

